# CX3CR1+ microglia/macrophages, activated T cells, and IFN-γ-driven stimulation confer age-dependent protective immunity in young-adult mice following β-coronavirus infection

**DOI:** 10.64898/2026.09.16.751725

**Authors:** Satavisha Ghosh, Bishal Hazra, Subhajit Das Sarma, Debanjana Chakravarty, Jayasri Das Sarma

## Abstract

Age-dependent variation in the immune response is a critical determinant of host susceptibility, disease severity, and long-term sequelae in coronavirus infections, a principle strikingly demonstrated by the COVID-19 pandemic caused by SARS-CoV-2. Although primarily pneumotropic, coronaviruses carry significant neurotropic potential, driving neurological complications whose severity is profoundly determined by host age. Elderly individuals and often children suffer disproportionately severe disease, whereas young adults relatively mount protective responses; yet the cellular and molecular determinants of this age-dependent neuroprotection remain poorly characterized. Using juvenile and young-adult C57BL/6 mice intracranially inoculated with β-coronavirus MHV-RSA59, a murine equivalent of human coronaviruses, we examined immune dynamics at days 5 (innate-acute), 7 (innate-to-adaptive transition), and 30 (chronic) post-infection. Young-adult mice exhibited only occasional demyelination, in stark contrast to the extensive demyelination observed across all spinal cord levels in juveniles. This differential outcome was attributable to enhanced age-dependent immune maturation, characterized by efficient T cell-microglia/macrophage crosstalk enabling effective viral control by day 7 post-infection. Young adults displayed greater glial activation, heightened cytokine release, CX3CR1+ microglia activation, and augmented CNS trafficking of CX3CR1+ and MHC II+ monocytes/macrophages, IFN-γ+ CD4+/CD8+ T cells, and CXCR3+ effector T cells relative to juveniles. Reduced naïve T cell frequencies and elevated effector/central memory T cell populations in cervical lymph nodes further indicate a robust adaptive memory response conferring long-term protection. Additionally, greater regulatory T cell accumulation in young adults facilitates timely suppression of excessive neuroinflammation as viral burden subsides. Together, these findings define the age-dependent immune landscape that underpins neuroprotection in β-coronavirus infection.

## Introduction

The age of the host is a fundamental determinant of immune system activation, maturation, and, consequently, of susceptibility to viral infection, disease severity, and the capacity to mount effective antiviral immunity. During infancy, childhood, and adolescence, both the innate and adaptive arms of the immune system undergo continuous developmental refinement encompassing the expansion of naïve lymphocyte pools, the progressive maturation of antigen-presenting cell function, and the gradual acquisition of immunological memory(1). These age-dependent immunological changes profoundly influence viral infectivity, the efficiency of viral clearance, and the risk of long-term sequelae following infection. Among the viral pathogens that are of a significant threat to humans, coronaviruses are of particular significance. Coronaviruses are classified into four genera based on their phylogenetic and genomic characteristics: Alphacoronavirus, Betacoronavirus, Gammacoronavirus, and Deltacoronavirus(2, 3). Among these, β-coronaviruses possess significant zoonotic potential and are responsible for severe respiratory diseases in humans(4, 5). Although primarily pneumotropic, accumulating evidence indicates that β-coronaviruses are also neurotropic, capable of invading the central nervous system (CNS) and causing neuroglial damage(6–11). Neurological complications associated with human coronaviruses (HCoVs) have been extensively documented across age groups, with particular impact on children and older adults(7, 8, 12–15). Indeed, SARS-CoV-2 infection and its long-term neurological sequelae, including febrile seizures, encephalitis, myelitis, meningitis, and demyelination, exemplify this age-dependent variation, underscoring the need for systematic mechanistic investigation across developmental age groups(16–18).

Although elderly individuals with comorbidities are disproportionately susceptible to severe disease, children generally exhibit milder symptoms. Systematic meta-analysis of infection fatality rates encompassing multiple countries demonstrated a striking exponential relationship between age and COVID-19 mortality risk, with extremely low infection fatality rates in children and young adults(19–21). However, this paradigm is challenged by reports of rare hyperinflammatory conditions such as Multisystem Inflammatory Syndrome in Children (MIS-C)(22–25), in which pediatric patients exhibited persistent inflammatory conditions, lymphopenia, and multiorgan involvement, including cardiac, renal, respiratory, hematological, gastrointestinal, or neurological(22, 25–28). The median age of MIS-C susceptibility was reported to be approximately 8-10 years (22, 25)across multiple large cohort studies, with the highest incidence reported among school-aged children, whereas infants and older adolescents accounted for comparatively fewer cases. The reduced incidence of MIS-C in older adolescents and young adults raises the possibility that developmental changes in immune regulation may be one factor that increases susceptibility; however, the mechanisms underlying this age distribution remain incompletely understood.

Cohort studies have generated multiple hypotheses regarding the immune cell populations that regulate coronavirus replication and spread(29–31). Some reports associate reduced CD4+ T cell responses and diminished IFN-γ production with severe outcomes in children(29), whereas others indicate that impaired adaptive immunity does not invariably lead to severe disease(30). These discrepancies underscore the need to characterize systemic innate and adaptive immune dynamics during both the acute phase and the post-viral clearance period. The lower disease burden in young adults relative to susceptible pediatric populations implies the existence of age-dependent protective mechanisms. However, most clinical studies are cross-sectional and lack longitudinal follow-up, limiting the ability to assess how patient age shapes disease severity and progression. Furthermore, the immune dynamics throughout the disease course and the cytokine milieu driving neuroinflammation remain largely unexplored. Murine models, which permit precise mechanistic dissection of immune variables across defined developmental stages, provide an indispensable experimental platform to address these knowledge gaps.

To investigate the immune environment in juvenile and young-adult hosts, a comparison not feasible in clinical settings, we used C57BL/6 mice from two defined age groups: 4-week-old mice, corresponding to approximately 9-10-year-old children, and 7-8-week-old mice, corresponding to 18-21-year-old young adults in humans(32). These mice were intracranially infected with the murine β-coronavirus mouse hepatitis virus (MHV-RSA59), a well-established murine equivalent of human coronaviruses. Infection of C57BL/6 mice with MHV-RSA59 provides a direct virus-induced experimental model for delineating the mechanisms of neuroglial damage, demyelination, and axonal loss(33–35). This model is therefore well-suited for investigating age-dependent variation in virus-induced neuropathogenesis, neuroinflammation, and demyelinating pathology across three defined stages of disease: day 5 (innate-acute phase), day 7 (innate-to-adaptive transition phase), and day 30 post-infection (chronic phase). Our initial examination of demyelination at day 30 p.i. revealed that microglia/macrophage-mediated demyelination was infrequent and markedly reduced in young-adult mice during the chronic stage. To elucidate the underlying mechanisms contributing to a tightly regulated and optimal immune response in young adults resulting in significantly reduced demyelination, we conducted a comprehensive investigation of the early stages of infection. This encompassed viral load analysis, glial cell activation, crosstalk between CNS-resident glial cells (microglia and astrocytes) and peripheral myeloid and lymphoid immune populations, immune cell maturation within the cervical lymph nodes (CLNs), and the cytokine milieu associated with the protective antiviral response in young-adult mice.

The present study demonstrates that neuroinflammation, driven by brain-resident glial cells and infiltrating peripheral immune populations, is optimally regulated in young-adult mice. During the early stages of infection, young-adult mice mounted a markedly stronger and more coordinated immune response than juvenile mice, effectively reducing viral load by day 7 p.i., the innate-to-adaptive transition stage. Robust immune cell maturation in the CLNs, consequent CNS infiltration, and extensive microglial and astrocyte activation at the initial disease stage collectively contributed to efficient viral control in the young-adult cohort. The principal immune populations mediating this protection were activated CX3CR1+ and MHC II+ macrophages, IFN-γ-producing CD4+/CD8+ T cells, and CXCR3+ effector T cells. Our previous studies indicate that CD4+ T cell-microglia/macrophage interaction not only contributes to a sustained CX3CR1 activation in microglia/macrophages during the acute neuroinflammatory phase required for viral clearance, but this bidirectional interaction also plays a crucial role in maintaining microglial homeostasis during chronic demyelination(36). The markedly greater CX3CR1 activation in the microglia of young adults, together with augmented infiltration of IFN-γ+ T cells and CXCR3+ effector T cells, supports efficient viral clearance. Conversely, impaired microglial CX3CR1 activation and suboptimal CD4+ T cell infiltration contribute to higher viral loads in juvenile mice(37, 38). Consistent with our earlier findings demonstrating that CX3CR1 expression is regulated by the interferon-stimulated gene ISG54 (encoding Ifit2) (37, 38)the present study reveals an age-dependent upregulation of Ifit2 that directly correlates with reduced viral burden in young-adult compared to juvenile mice. Additionally, greater regulatory T cell (Treg) infiltration and enhanced TGF-β production in young-adult mice facilitate timely suppression of exacerbated neuroinflammation and prevent excessive immune activation as viral burden declines. Finally, the lower pool of naïve T cells and the significantly higher proportion of effector and central memory T cells in the CLNs of young-adult mice compared to juveniles indicate robust age-dependent immune maturation and a long-term protective immune response.

## Results

### The percentage area of demyelination was significantly lower in young adult mice in the sampled spinal cord sections at the chronic stage of infection

To evaluate the impact of age on long-term disease outcomes, 4-week-old and 7-8-week-old C57BL/6 mice were intracranially injected with MHV-RSA59 and sacrificed on day 30 post-infection. This time point represents the chronic stage of the disease, when replicating infectious virus particles are typically no longer prevalent in the CNS. Demyelinating lesions (indicated by dotted lines) were visible within the white matter tracts of the spinal cord. While the 4-week-old juvenile mice exhibited extensive demyelination in both the dorsal and ventral columns, the young adult mice displayed occasional and only sparse demyelination in these regions (Fig. 1A). Specific details regarding the myelin staining protocol and spatial sectioning intervals along the spinal cord are detailed within the Materials and Methods section.

**Fig 1:**
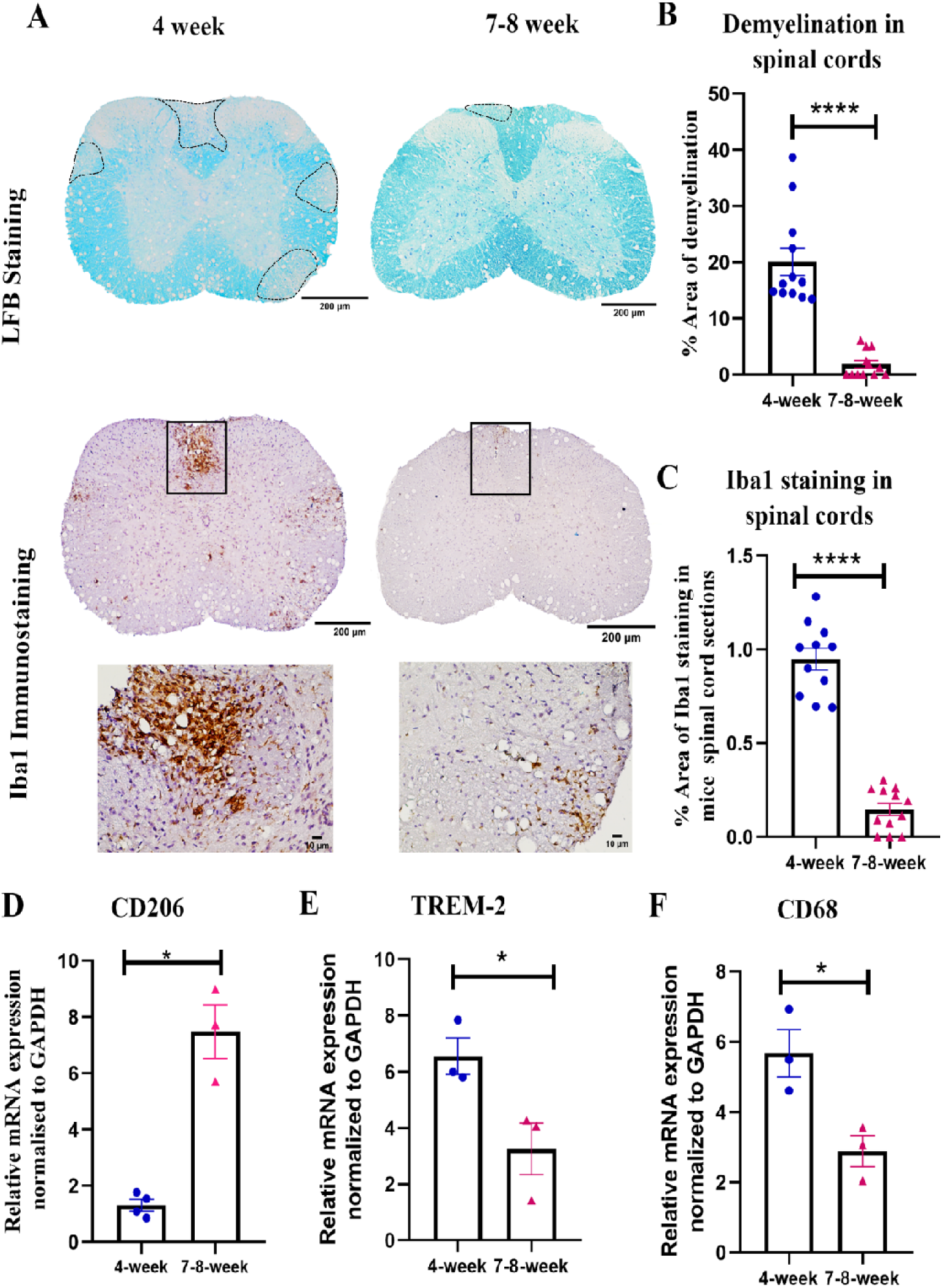
At the chronic phase, day 30 p.i., demyelination pathology was significantly reduced and rarely observed in young adult mice: **A)** On day 30 p.i., spinal cord cross-sections of MHV-RSA59-infected 4-week and 7-8-week-old mice were analyzed for the presence of demyelinating plaques by LFB staining. Black dotted lines represent demyelinating plaques in the spinal cord’s white matter. Activated microglia/macrophages around the demyelinating plaques were immunostained by Iba1. The black boxed region in respective spinal cord sections captured at higher magnification in insets (40X). Scale bar 200µm, 10µm. **B)** Quantification of the percentage area of demyelination. **C)** Quantification of Iba1 immunostaining. **D-F)** Abundance of *CD206*, *Trem2*, and *CD68* transcripts was determined in the spinal cords by qRT-PCR. The mRNA levels were normalized to *Gapdh*, which was used as the housekeeping control. The fold changes were compared to the mock-infected control group. Results were expressed as mean ± SEM. The shown histopathological studies are representative of two separate experiments, and qRT-PCR study is representative of one independent experiment. (n=3-6 per age group, 1-2 technical replicates have been plotted from each mouse for histopathological studies). *Asterisk represents statistical significance calculated using an unpaired Student’s t-test; P<0.05 was considered significant. ****P<0.001.

Accordingly, quantification of the percentage area of demyelination revealed significantly less demyelinating lesion burden in the young adult mice, showing that at day 30 p.i., the percentage area of demyelination was lower in young adult mice in the sampled spinal cord sections (Fig. 1B). In mice from both age groups, gray matter inflammation was largely resolved by this chronic stage.

The areas of white matter demyelination were next examined for the presence of activated phagocytic microglia/macrophages (Fig. 1A, black boxes) in spinal cord serial sections, as these immune cells play a crucial role in engulfing myelin and actively executing myelin stripping(33). Quantification of Iba1 (Ionized calcium-binding adaptor protein-1) staining intensity revealed a significantly higher persistence of activated microglia/macrophages within the demyelinating lesions of the spinal cord white matter in the juvenile 4-week-old mice. Conversely, the young adult group showed significantly fewer Iba1-immunoreactive activated microglia/macrophages surrounding the demyelinating lesions (Fig. 1C).

Furthermore, RNA isolated from day 30 spinal cord tissues was subjected to qRT-PCR to assess microglial activation profiles at the transcript level. These data demonstrated a significantly higher expression of CD206, a marker associated with anti-inflammatory, reparative microglia (Fig. 1D), in the spinal cords of young adult mice compared to juveniles. Concurrently, the expression of microglial phagocytic markers such as TREM-2 (Fig. 1E) and CD68 (Fig. 1F) was significantly lower in the young adult cohorts. The significant reduction in Iba1 immunoreactivity, along with shifts in microglial activation markers at the transcript level (CD206(up), TREM-2(Down), CD68(Down)) at day 30 (Fig. 1C-F), were highly consistent with a reduced demyelinating lesion burden in young adults.

### Young adult mice cleared infectious viral particles more efficiently by day 7 p.i., consistent with a more effective antiviral immune response

To understand the pathophysiological processes that culminate in enhanced chronic demyelination in juvenile mice, the disease pathology at the initial stages of infection was investigated next, focusing on the acute-innate phase (day 5 p.i.) and the innate-adaptive transition phase (day 7 p.i.) when the replicating viruses are actively present in the system. Mice from both age groups were monitored daily for clinical symptoms following infection (Fig. 2). Longitudinal analysis of body weight revealed comparable weight changes between the experimental cohorts across both age groups, displaying a consistent pattern of weight loss that continued until days 10-11 p.i. (Fig. 2A). Mice from both groups displayed an initial average clinical score of 1.0-1.5, characterized by ruffled fur and a prominent hunchback phenotype. However, after day 8 p.i., the clinical scores in juvenile mice worsened significantly until day 13 p.i., before showing gradual signs of recovery. Occasionally, some juvenile mice exhibited severe disease scores of 2.0-2.5, marked by partial or complete hind limb paralysis, whereas such high clinical scores were notably absent in the young adult group (Fig. 2B). Disease severity was correlated with viral load analysis by assessing the number of infectious viral particles in the mice brains on days 3, 5, and 7 p.i. using a routine standard viral plaque assay. The viral load was reported to be equivalent between the two age groups on days 3 and 5 p.i.; however, on day 7 p.i., infectious viral titers were significantly reduced in the young adult mouse cohort (Fig. 2C). To complement this, the expression of the viral N gene was assessed at the transcript level via qRT-PCR, which revealed comparable levels of viral N gene RNA in mice from both age groups during early timepoints, including days 3, 5, and 7 p.i. (Fig. 2D).

**Fig 2:**
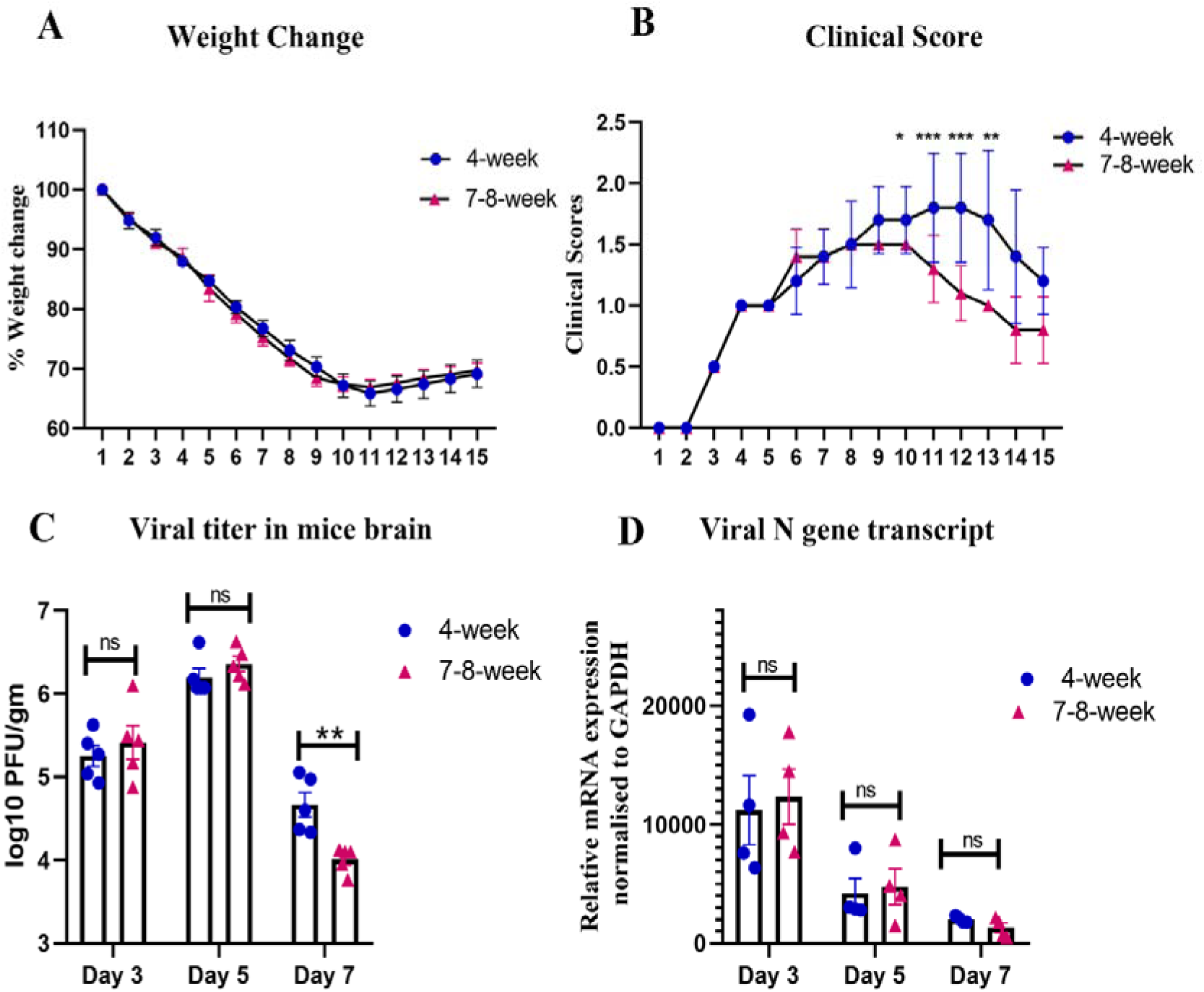
Despite similar percent weight change between juvenile and young adult mice, clinical scores were lower in the young adult mice at the innate-adaptive transition phase, with significant reduction of viral load in the CNS post-infection with MHV-RSA59: 4-week and 7-8-week-old mice were infected with MHV-RSA59 and monitored daily for **A)** weight change and **B)** development of clinical score. Clinical scores were assigned by a relative scale of 0-4, (detailed in material and methods section). **C)** viral titers in brains on days 3, 5, and 7 p.i. **D)** The abundance of viral *N-gene* transcript was determined in brain lysates on days 3, 5 and 7 p.i. by qRT-PCR. The *N-gene* mRNA levels were normalized to *Gapdh*, which was used as the housekeeping control. The fold changes were compared to the mock-infected control group. Results were expressed as mean ± SEM. The shown weight change, clinical score study, viral titer analysis and qRT-PCR study are representative of one independent experiment. (n=4-5 per age group). * Asterisk represents statistical significance calculated using Two-Way ANOVA analysis. P<0.05 was considered significant, **P<0.01, ***P<0.001.

Although identical volumes of viral inoculum were administered to both age groups, despite differences in brain mass, the equivalent level of infectious viral particles and comparable viral N gene transcript levels at the initial stages of the disease (Fig. 2C, D) indicate that comparable levels of viral replication were established in both age groups after infection. This important benchmark suggests that the reduced infectious viral titers at day 7 (Fig. 2C) reflect genuine differences in viral clearance kinetics rather than the effects of the initial inoculum. Cumulatively, these data indicate that, despite comparable early viral establishment, as evidenced by equivalent viral titers and N gene transcripts at days 3, 5, and 7 p.i., young adult mice cleared infectious viral particles more efficiently by day 7 p.i., consistent with a more effective antiviral immune response in this age group.

### Encephalitis was heightened in the brain of young adult mice at the early stages of infection, with a greater activation of brain-resident glial cells

Given that young adult mice exhibited accelerated clearance of infectious virus by day 7, it was imperative to investigate the kinetics of neuroinflammation. H&E-stained brain sections from MHV-RSA59-infected juvenile and young adult mice revealed enhanced acute encephalitis characterized by perivascular lymphocytic cuff formation (indicated by arrow) and microglial nodule formation (indicated by arrow) at both the acute-innate phase (day 5 p.i.) and the innate-adaptive transition phase (day 7 p.i.) (Fig. 3A and Fig. 4A).

**Fig 3:**
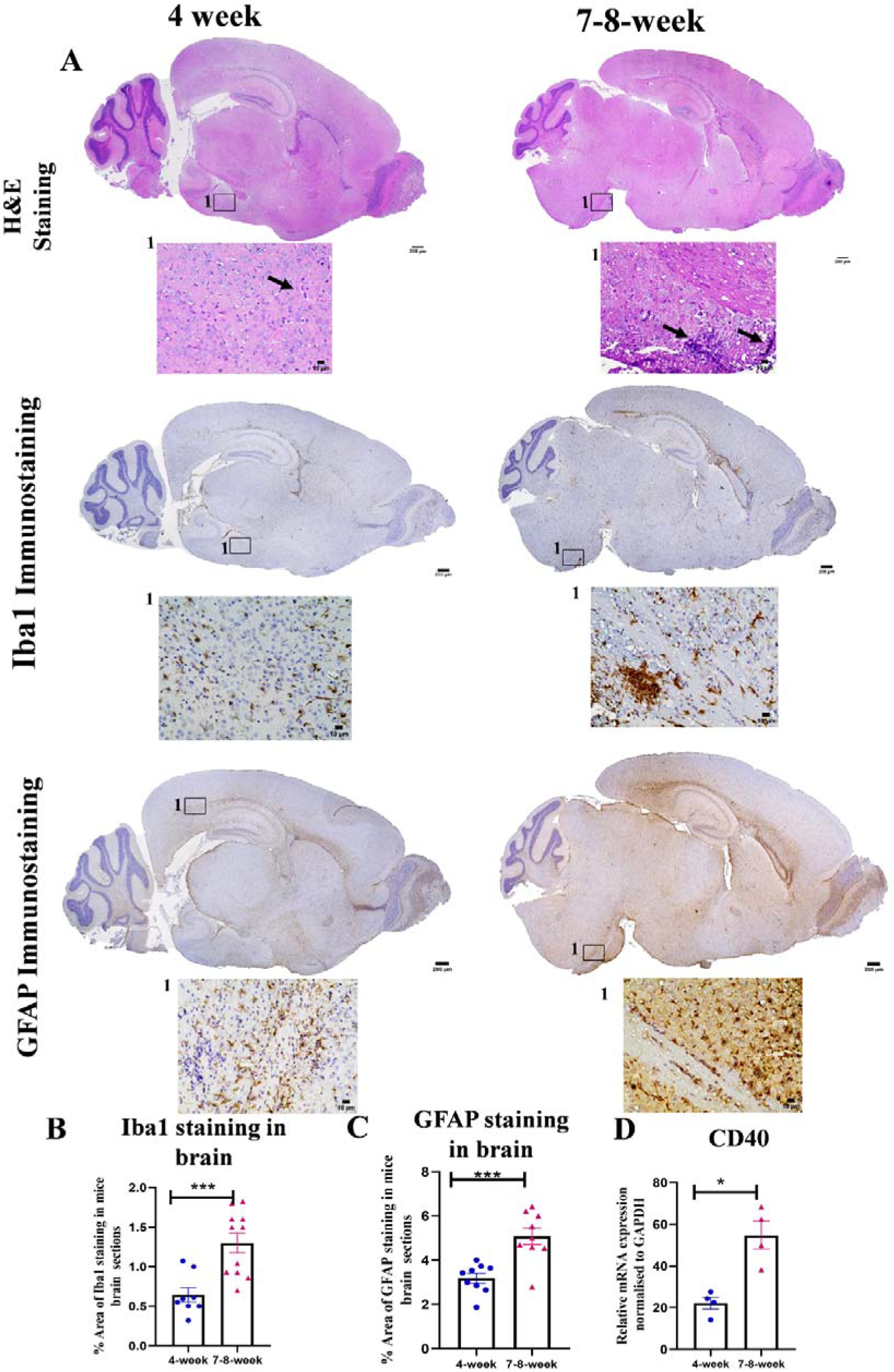
The young adult mice exhibited a higher microglial and astrocyte activation at the acute phase of neuroinflammation: **A)** On day 5 p.i., mid-sagittal sections of brains from MHV-RSA59-infected 4-week and 7-8-week-old mice were stained with H&E and immunostained with Iba1 and GFAP. The black boxed region in brain sections captured at higher magnification in insets (40X) below the respective brain sections. Scale bar 200µm, 10µm. **B)** Quantification of Iba1 immunostaining. **C)** Quantification of GFAP immunostaining. **D)** The abundance of *CD40* at the transcript level was determined in brain lysates by qRT-PCR. The mRNA levels were normalized to *Gapdh*, which was used as the housekeeping control. The fold changes were compared to the mock-infected control group. Results were expressed as mean ± SEM. The shown histopathological and qRT-PCR studies are representative of one independent experiment. (n=4-6 per age group, 1-2 technical replicates per mouse have been plotted for histopathological experiments) *Asterisk represents statistical significance calculated using an unpaired Student’s t-test; P<0.05 was considered significant. ***P<0.001.

**Fig 4:**
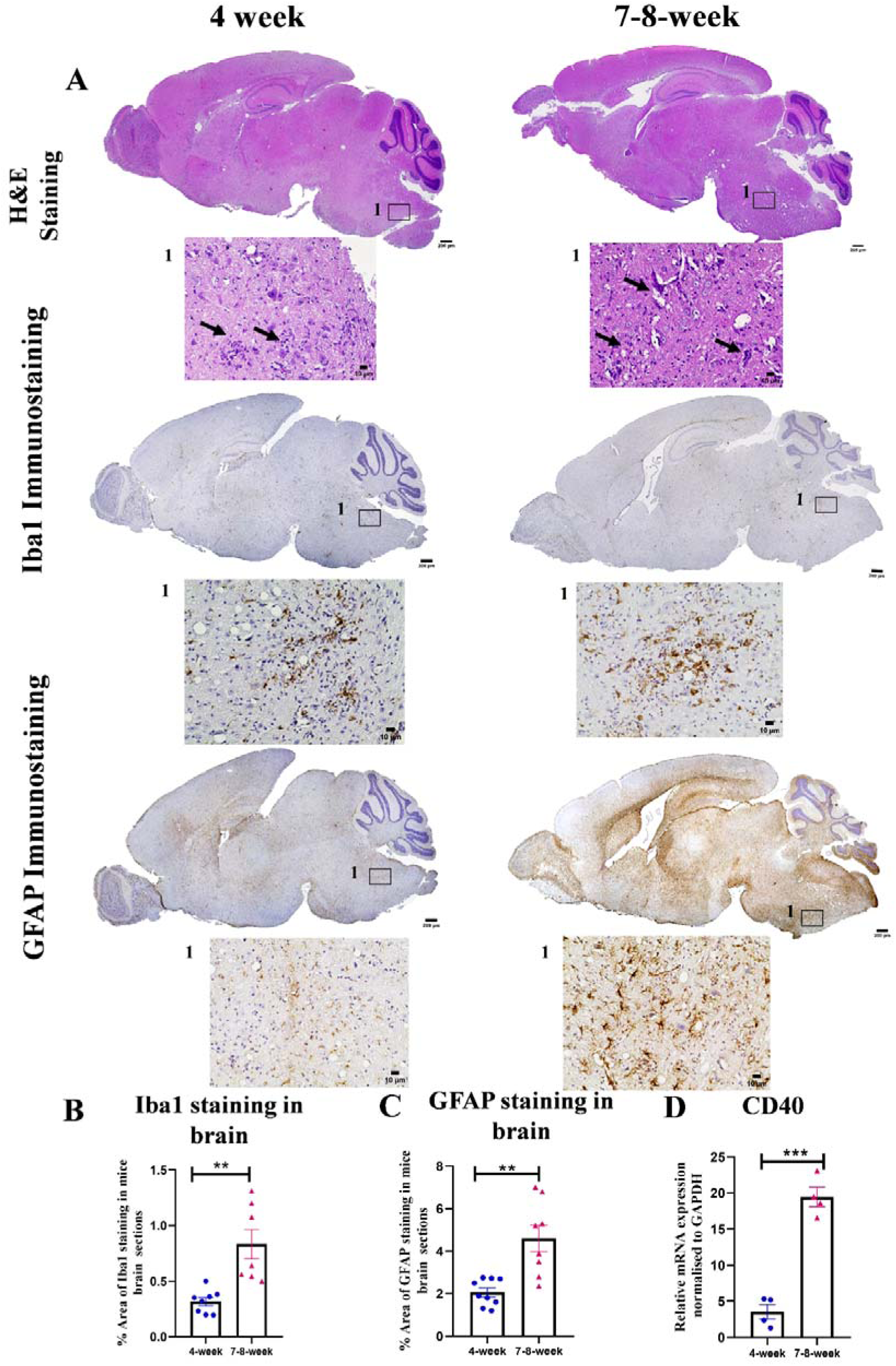
At the innate-adaptive transition phase, the young adult mice also exhibited higher microglial and astrocyte activation: **A)** On day 7 p.i., mid-sagittal brain sections from MHV-RSA59 infected 4-week and 7-8-week-old mice were stained with H&E and immunostained with Iba1 and GFAP. The black boxed region in brain sections captured at higher magnification in insets (40X) below the respective brain sections. Scale bar 200µm, 10µm. **B)** Quantification of Iba1 immunostaining. **C)** Quantification of GFAP immunostaining. **D)** The abundance of *CD40* at the transcript level was determined in brain lysates by qRT-PCR. The mRNA levels were normalized to *Gapdh*, which was used as the housekeeping control. The fold changes were compared to the mock-infected control group. Results were expressed as mean ± SEM. The shown histopathological and qRT-PCR studies are representative of one independent experiment. (n=4-5 per age group, 1-2 technical replicates per mouse have been plotted for histopathological experiments) *Asterisk represents statistical significance calculated using an unpaired Student’s t-test; P<0.05 was considered significant. **P<0.01, ***P<0.001.

Corresponding serial brain sections were immunostained with anti-Iba1 to assess microglial activation dynamics (Fig. 3A and Fig. 4A). Quantification of Iba1 staining intensity revealed a significantly higher prevalence of activated microglia/macrophages in the brains of young adult mice when compared to the juvenile groups at these early timepoints (Fig. 3B and Fig. 4B). Parallel immunostaining experiments using anti-GFAP (Glial fibrillary acidic protein) to locate reactive astrocytes yielded similar patterns (Fig. 3A and Fig. 4A), with quantification demonstrating significantly higher astrocyte activation intensity in the brains of young adult mice at both day 5 and day 7 p.i. (Fig. 3C and 4C).

Furthermore, RNA extracted from day 5 and day 7 brain tissues was analyzed by qRT-PCR for CD40, a critical co-stimulatory activation marker expressed on antigen-presenting microglia and macrophages. This analysis indicated a significantly higher expression of CD40 transcripts in the young adult mice when compared to the juvenile mice at both examined timepoints (Fig. 3D and 4D). Cumulatively, these findings indicate that, with advancing age into young adulthood, a more robust, protective early response by activated glial cells correlates with reduced viral invasion and accelerated clearance.

### The onset of demyelination occurs during the acute-innate phase (day 5 p.i.) in juvenile mice, while myelitis is evident across both age groups

H&E-stained spinal cord cross-sections from infected juvenile and young adult mice revealed pronounced acute myelitis (indicated by arrow), with white and gray matter inflammation characterized by prominent microglial nodule formation at day 5 p.i. in both cohorts (Fig. 5A). However, when evaluating the precise onset of demyelination pathology at this acute-innate stage, while infectious viral particles are actively replicating throughout the CNS, clear differences were observed. Small, distinct demyelinating lesions (indicated by dotted arrows) were significantly visible in the spinal cord white matter tracts of the juvenile mice. In contrast, no significant focal demyelinating plaques were observed in the young adult group at this stage (Fig. 5A).

**Fig 5:**
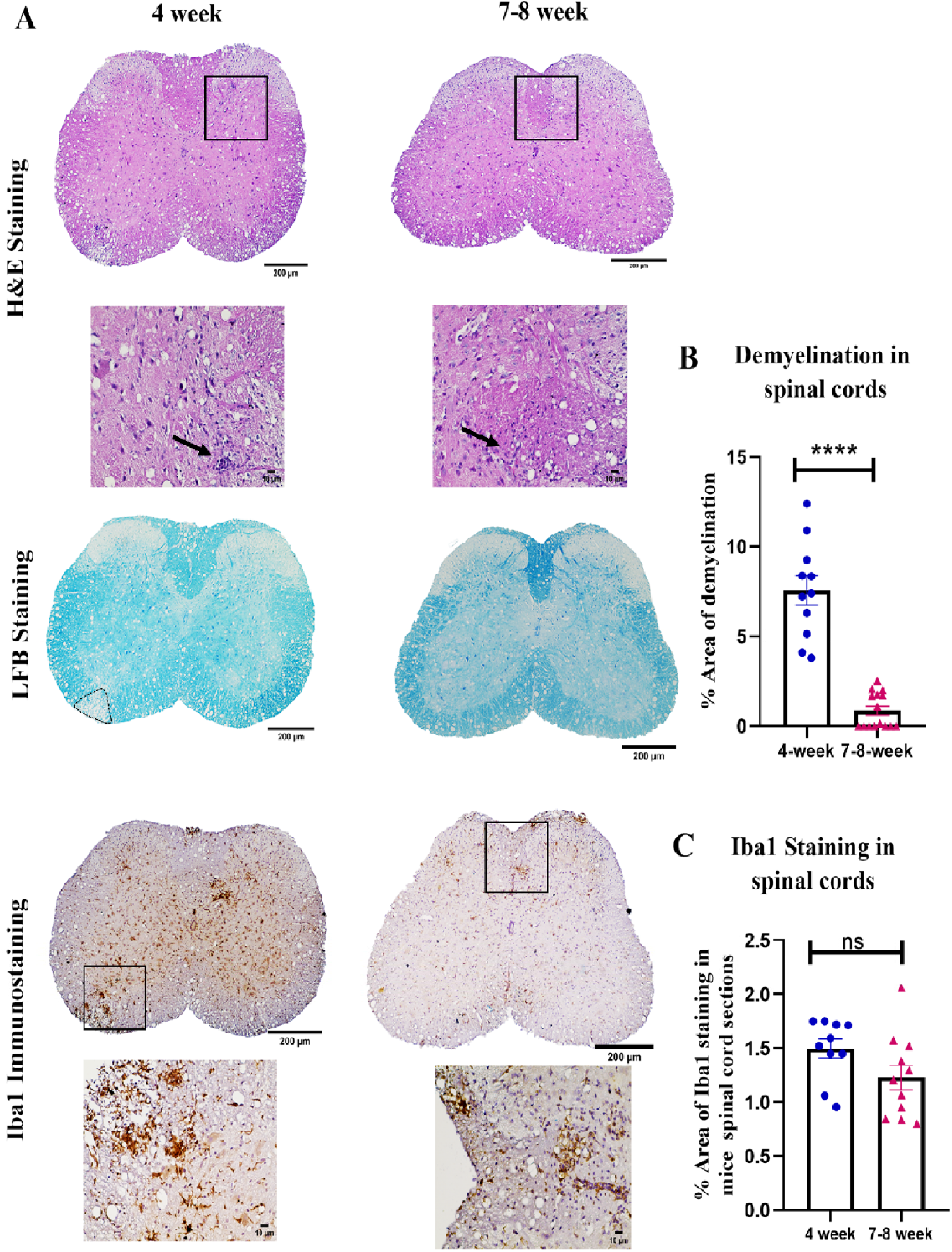
The young adult mice exhibited an occasional demyelination onset on day 5 p.i.: **A)** On day 5 p.i., spinal cord sections of MHV-RSA59 infected 4-week and 7-8-week-old mice were analyzed for the presence of inflammatory lesions and myelitis by H&E staining, demyelinating plaques by LFB staining. Black dotted lines represent demyelinating plaques in the spinal cord’s white matter. Activated microglia/macrophages in the gray and white matter regions by Iba1 immunostaining. The black boxed region in respective spinal cord sections captured at higher magnification in insets (40X). Scale bar 200µm, 10µm. **B)** Quantification of the percentage area of demyelination. **C)** Quantification of Iba1 immunostaining. The shown histopathological studies are representative of two separate experiments. (n=5 per age group, 2-3 technical replicates per mouse have been plotted). *Asterisk represents statistical significance calculated using an unpaired Student’s t-test; P<0.05 was considered significant. ****P<0.0001.

While all juvenile mice exhibited distinct demyelinating plaques in the dorsal and ventral columns, young adult mice showed only sparse, isolated demyelination restricted to the dorsal column. Quantification of the percentage area of demyelination onset at this acute stage confirmed a significantly greater extent of early demyelination development in the juvenile group, whereas demyelination area was almost negligible in the young adults (Fig. 5B).

Furthermore, the areas of white matter demyelination and gray and white matter inflammation were examined for the presence of Iba1+ activated microglia/macrophages in serial spinal cord sections (Fig. 5A, black boxes). Interestingly, quantification of the spinal cord Iba1 staining intensity at day 5 p.i. revealed comparable overall activation levels of microglia/macrophages within the spinal cord tissues of both age groups (Fig. 5C), indicating that the early protection against demyelination onset in adults is not due to a localized absence of microglia/macrophage activation.

### Whole-brain analyses show higher CNS-associated myeloid cell numbers and activation in young adult mice on day 5 p.i

To precisely define which immune cells constitute the early inflammatory response, flow cytometric analysis was performed on whole-brain lysates to enumerate absolute cell numbers of peripheral leukocytes and brain-resident microglia during the acute-innate stage (day 5 p.i.; Fig. 6) and the innate-adaptive transition stage (day 7 p.i.; Fig. S2). The detailed gating strategies for these flow cytometry experiments are outlined in the Materials and Methods section. The mock-infected (MI) control mice confirmed that no significant differences existed in the populations of brain-resident microglia or peripheral leukocytes between MI juveniles and MI young adult mice (Fig. S1A, B and C, D).

**Fig 6:**
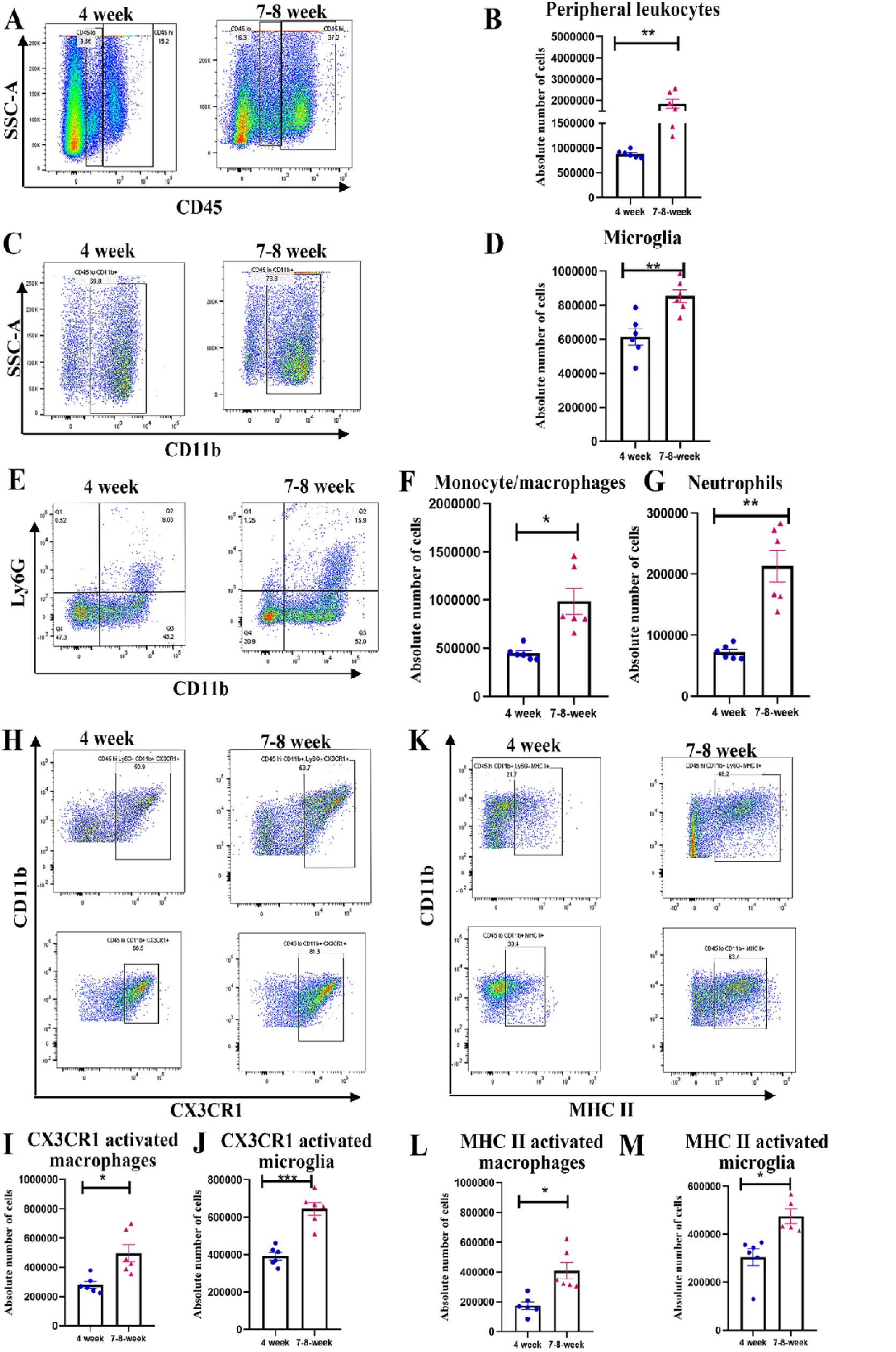
The infiltration and activation of peripheral monocyte/macrophages, as well as the numbers and activation of brain resident microglia, were significantly higher in young adult mice. On day 5 p.i., brains from MHV-RSA59-infected 4-week and 7-8-week-old C57BL/6 mice were harvested for flow cytometry analysis and stained for CD45, CD11b, Ly6G, CX3CR1, and MHC II. Primary gating was performed on live cells, followed by singlets and CD45 hi. **A)** Representative flow cytometry dot plots showing percentages of overall CD45hi and CD45lo cell populations after gating on live cells followed by singlets. **B)** Graphical representation of the absolute cell numbers of peripheral leukocytes. **C)** Representative flow cytometry plots showing CD45lo CD11b+ (brain resident microglia). **D)** Graphical representation of the absolute numbers of brain resident microglia. **E)** Representative flow cytometry dot plots showing percentages of CD45hi CD11b+ Ly6G-(peripheral monocyte/macrophages) and CD45hi CD11b+ Ly6G+ (neutrophils). **F)** Graphical representation of the absolute cell numbers of infiltrating monocyte/macrophages **G)** neutrophils. Representative flow cytometry dot plots showing CD45hi CD11b+ Ly6G-cells and CD45lo CD11b+ cells expressing activation markers **H)** CX3CR1 **K)** MHC-II. Graphical representations of the absolute cell numbers have been shown in **I) J) L)** and **M),** respectively. Results were expressed as mean ± SEM. The shown flow cytometry study is representative of one independent experiment. (n=5-6 per age group) *Asterisk represents statistical significance calculated using unpaired Student’s t-test, P<0.05 was considered significant, **P<0.01, ***P<0.001.

Following viral infection, whole-brain analyses show higher CNS-associated myeloid cell numbers and activation in young adult mice on day 5 post-infection. At this acute stage, infected young adult mice exhibited significantly greater total numbers of CNS-associated peripheral leukocytes compared to juveniles (Fig. 6A, B). Similarly, quantitative analysis of the brain-resident microglial compartment revealed a significantly larger absolute population of microglia in young adults on day 5 p.i. relative to juvenile cohorts (Fig. 6C, D).

Further dissection of specific myeloid subsets revealed no baseline variations in monocyte/macrophage or neutrophil numbers between mock-infected control groups (Fig. S1E, F, G). Upon infection, however, both monocyte/macrophage (Fig. 6E, F) and neutrophil (Fig. 6E, G) numbers were significantly higher in the young adult brain compared to juveniles.

The activation status of these cell types was evaluated by measuring the surface expression of major histocompatibility complex class II (MHC II) and the CX3C chemokine receptor 1 (CX3CR1). Comparing the infected age groups revealed a significantly higher accumulation of CX3CR1-activated macrophages (Fig. 6H, I) and MHC II-activated macrophages (Fig. 6K, L) in the young adult group. The abundance of CX3CR1-activated microglia (Fig. 6H, J) and MHC II-activated microglia (Fig. 6K, M) was similarly increased in young adults at this acute stage. Conversely, by the transition phase at day 7 p.i., the flow cytometric data revealed an overall decline in myeloid cell accumulation, with the numbers of peripheral monocytes/macrophages and neutrophils becoming statistically comparable between the two age groups (Fig. S2). Cumulatively, these data indicate that young adult mice mount a highly robust early innate host response upon MHV-RSA59 infection, characterized by the rapid accumulation of activated myeloid cell subsets and by the greater persistence of activated brain-resident microglia.

### Infiltration of lymphoid cell lineages, effector T cells, and IFN-γ-expressing cells was significantly higher in young adult mice

Next, the accumulation of lymphoid lineage cells within the CNS was evaluated. Mock-infected mice from both age groups displayed no alterations or age-dependent deviations in baseline lymphoid cell numbers (Fig. S1H-M). Following infection, the absolute numbers of CD4+ T cells (Fig. 7A, C), IFN-γ-expressing CD4+ T cells (Fig. 7B, D), CD8+ T cells (Fig. 7E, G), IFN-γ-expressing CD8+ T cells (Fig. 7F, H), and CXCR3+ effector CD8+ T cells (Fig. 7J, M) were significantly higher in the young adult group compared to juveniles at the acute stage (day 5 p.i.). At this early time point, the accumulation of CXCR3+ effector CD4+ T cells (Fig. 7I, L) and NKT cells (Fig. 7K, N) remained equivalent between the two infected cohorts.

**Fig 7:**
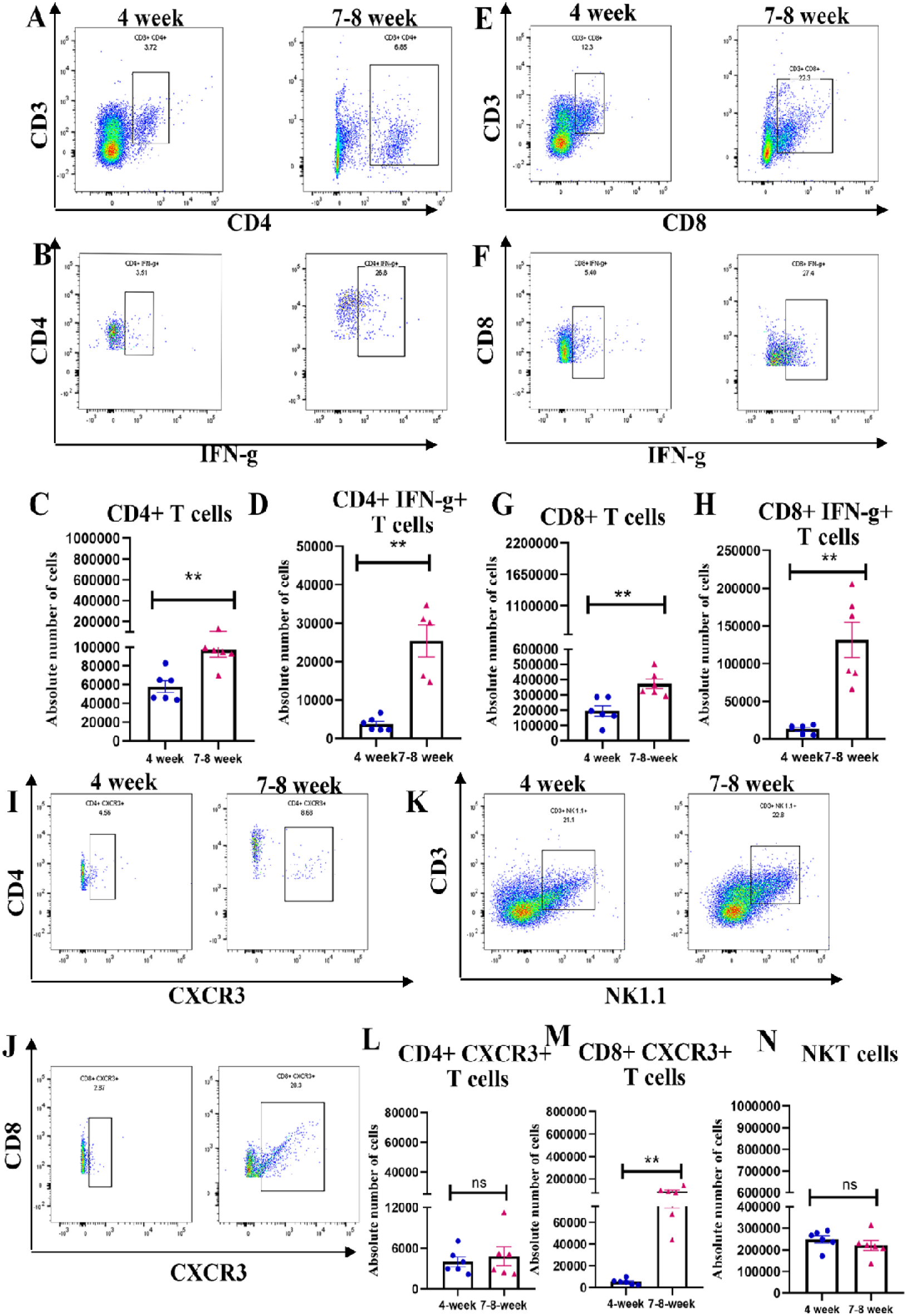
The infiltration of the cells of the lymphoid lineage was significantly higher in young adult mice at the acute phase of neuroinflammation. On day 5 p.i., brains from MHV-RSA59-infected 4-week and 7-8-week-old C57BL/6 mice were harvested for flow cytometry analysis and stained for CD45, CD3, CD4, CD8, NK1.1, CXCR3, and IFN-γ. Primary gating was performed on live cells, followed by singlets and CD45 hi. Representative flow cytometry dot plots showing percentages of **A)** CD3+ CD4+ T cells, **B)** CD3+ CD4+ IFN-γ+ T cells. Graphical representation of the absolute cell numbers of **C)** infiltrating CD4+ T cells, **D)** CD4+ IFN-γ+ T cells. Representative flow cytometry dot plots showing percentages of **E)** CD3+ CD8+ T cells, **F)** CD3+ CD8+ IFN-γ+ T cells. Graphical representation of the absolute cell numbers of **G)** infiltrating CD8+ T cells, **H)** CD8+ IFN-γ+ T cells. Representative flow cytometry dot plots showing percentages of **I)** CD3+ CD4+ CXCR3+ T cells, **J)** CD3+ CD8+ CXCR3+ T cells. Graphical representation of the absolute cell numbers of **L)** CXCR3+ CD4+ T cells, **M)** CXCR3+ CD8+ T cells. Representative flow cytometry dot plots showing percentages of **K)** CD3+ NK1.1+ (NKT cells). **N)** Graphical representation of the absolute cell numbers of infiltrating NKT cells. Results were expressed as mean ± SEM. The shown flow cytometry study is representative of one independent experiment. (n=5-6 per age group) *Asterisk represents statistical significance calculated using unpaired Student’s t-test, P<0.05 was considered significant, **P<0.01.

Notably, the infiltration of these lymphoid immune cell populations increased markedly as infection progressed into the innate-adaptive transition stage, reaching peak levels by day 7 p.i. At this transition phase, the accumulation of CD4+ T cells (Fig. 8A, C), IFN-γ-expressing CD4+ T cells (Fig. 8B, D), IFN-γ-expressing CD8+ T cells (Fig. 8F, H), CXCR3+ effector CD4+ T cells (Fig. 8I, L), CXCR3+ effector CD8+ T cells (Fig. 8J, M), and NKT cells (Fig. 8K, N) was significantly higher in the infected young adult mice compared to the juvenile group. Interestingly, the total number of CD8+ T cells (Fig. 8E, G) became comparable between the two age groups by day 7 p.i. Overall, these immune kinetics data demonstrate that lymphoid cell recruitment and functional activation steadily enhance during the innate-adaptive transition stage in young adults relative to juveniles.

**Fig 8:**
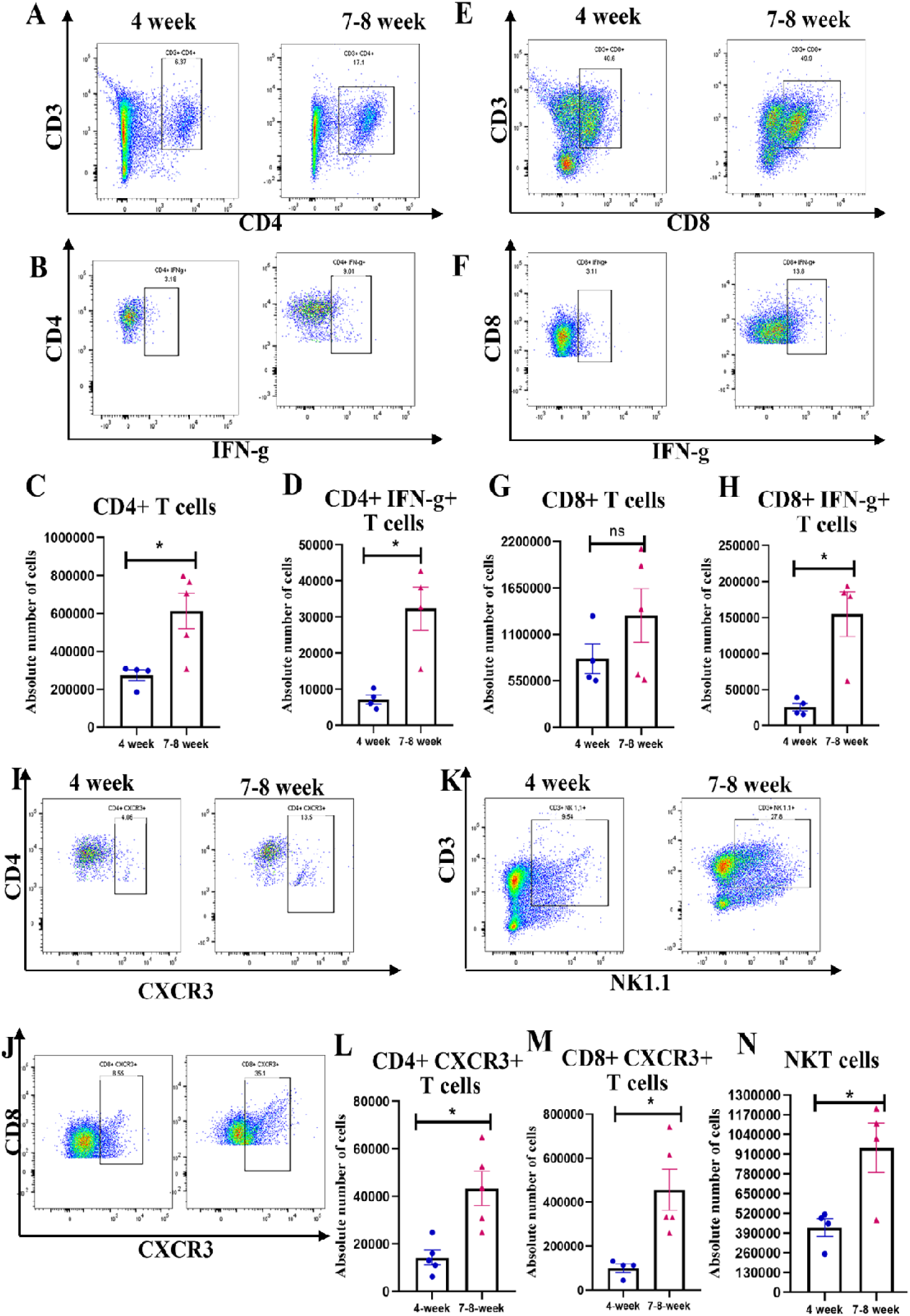
The infiltration of lymphoid lineage cells reached peak levels on day 7 p.i., and was significantly higher in young adult mice. On day 7 p.i., brains from MHV-RSA59-infected 4-week and 7-8-week-old C57BL/6 mice were harvested for flow cytometry analysis and stained for CD45, CD3, CD4, CD8, NK1.1, CXCR3, and IFN-γ. Primary gating was performed on live cells, followed by singlets and CD45 hi. Representative flow cytometry dot plots showing percentages of **A)** CD3+ CD4+ T cells, **B)** CD3+ CD4+ IFN-γ+ T cells. Graphical representation of the absolute cell numbers of **C)** infiltrating CD4+ T cells, **D)** CD4+ IFN-γ+ T cells. Representative flow cytometry dot plots showing percentages of **E)** CD3+ CD8+ T cells, **F)** CD3+ CD8+ IFN-γ+ T cells. Graphical representation of the absolute cell numbers of **G)** infiltrating CD8+ T cells, **H)** CD8+ IFN-γ+ T cells. Representative flow cytometry dot plots showing percentages of **I)** CD3+ CD4+ CXCR3+ T cells, **J)** CD3+ CD8+ CXCR3+ T cells. Graphical representation of the absolute cell numbers of **L)** CXCR3+ CD4+ T cells, **M)** CXCR3+ CD8+ T cells. Representative flow cytometry dot plots showing percentages of **K)** CD3+ NK1.1+ (NKT cells). **N)** Graphical representation of the absolute cell numbers of infiltrating NKT cells. Results were expressed as mean ± SEM. The shown flow cytometry study is representative of one independent experiment. (n=4-5 per age group) *Asterisk represents statistical significance calculated using an unpaired Student’s t-test; P<0.05 was considered significant.

### CLN T cell activation and indicators of memory T cells were higher in young adults at days 5-7 p.i., suggesting accelerated T cell priming

Because young adult mice exhibited a heightened accumulation of CD4+ and CD8+ T cells within the CNS, T cell priming and the kinetics of differentiation of memory T cell subsets were subsequently evaluated in the draining cervical lymph nodes (CLNs). Following infection, the total CD4+ T cell population was significantly higher in the CLNs of young adult mice at the acute stage, day 5 p.i. (Fig. 9A, C), before balancing out to comparable levels between the two age groups at day 7 p.i. transition stage (Fig. 10A, C). However, total IFN-γ-expressing CD4+ T cells (Fig. 9B, D and Fig. 10B, D), CD8+ T cells (Fig. 9E, G and Fig. 10E, G), and IFN-γ-expressing CD8+ T cells (Fig. 9F, H and Fig. 10F, H) were significantly more abundant in the young adult CLNs at both examined stages of infection compared to juvenile mice.

**Fig 9:**
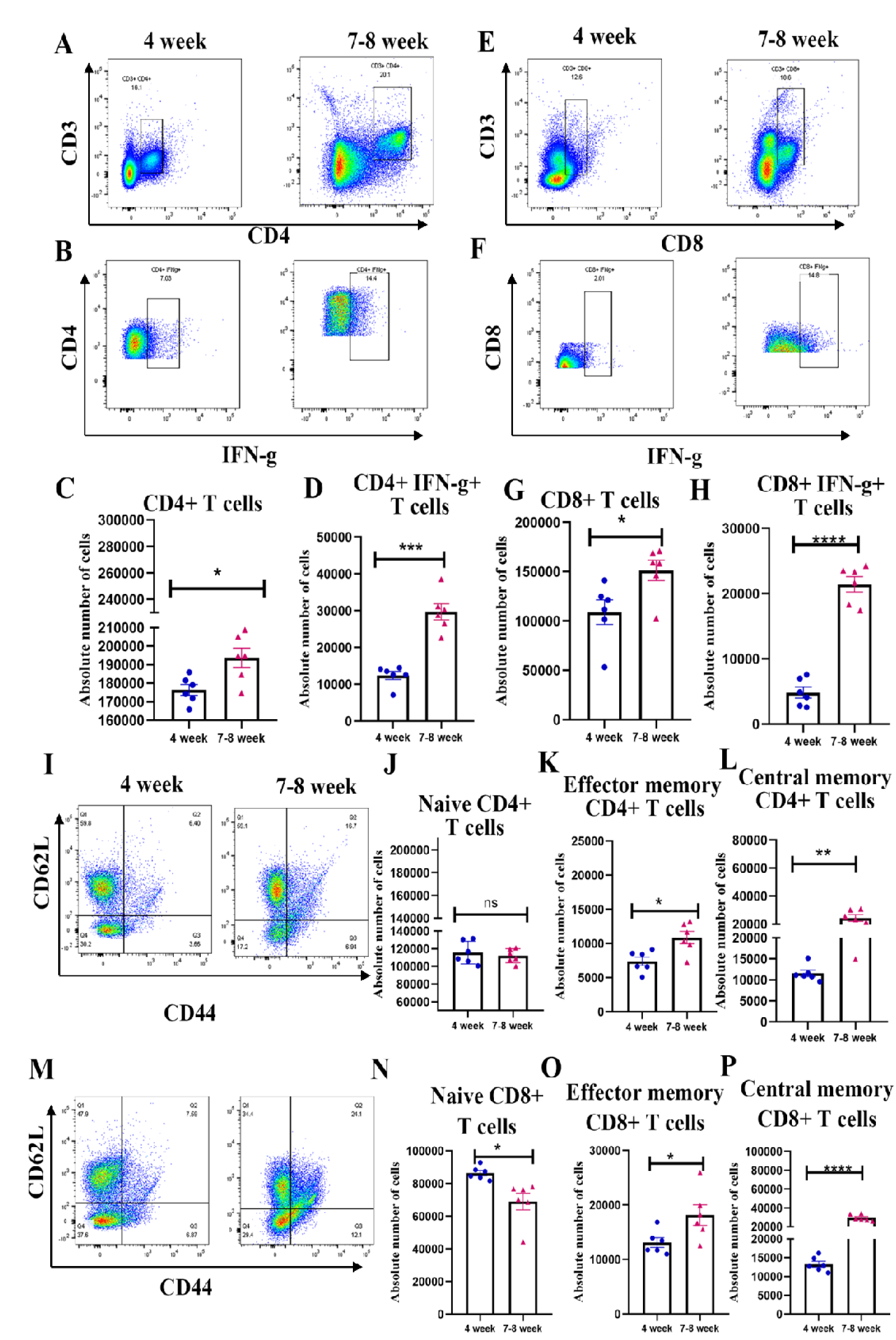
The priming activation and maturation of T cells were significantly enhanced in the young adult cohort at the acute phase of neuroinflammation. On day 5 p.i., CLNs from MHV-RSA59-infected 4-week and 7-8-week-old C57BL/6 mice were harvested for flow cytometry analysis and stained for CD3, CD4, CD8, IFN-γ, CD62L, and CD44. Primary gating was performed on live cells, followed by singlets. Representative flow cytometry dot plots showing percentages of **A)** CD3+ CD4+ T cells, **B)** CD3+ CD4+ IFN-γ+ T cells. Graphical representation of the absolute cell numbers of **C)** CD4+ T cells, **D)** IFN-γ expressing CD4+ T cells. Representative flow cytometry dot plots showing percentages of **E)** CD3+ CD8+ T cells, **F)** CD3+ CD8+ IFN-γ+ T cells. Graphical representation of the absolute cell numbers of **G)** CD8+ T cells, **H)** IFN-γ expressing CD8+ T cells. **I)** Representative flow cytometry dot plots showing percentages of CD4+ CD62L+ CD44-(Naïve CD4+ T cells), CD4+ CD62L-CD44+ (Effector memory CD4+ T cells), CD4+ CD62L+ CD44+ (Central memory CD4+ T cells). Graphical representation of the absolute cell numbers of **J)** Naïve, **K)** effector memory, and **L)** central memory CD4+ T cells. **M)** Representative flow cytometry dot plots showing percentages of CD8+ CD62L+ CD44-(Naïve CD8+ T cells), CD8+ CD62L-CD44+ (Effector memory CD8+ T cells), CD8+ CD62L+ CD44+ (Central memory CD8+ T cells). Graphical representation of the absolute cell numbers of **N)** Naïve, **O)** effector memory, and **P)** central memory CD8+ T cells. Results were expressed as mean ± SEM. The shown flow cytometry study is representative of one independent experiment. (n=6 per age group) *Asterisk represents statistical significance calculated using unpaired Student’s t-test, P<0.05 was considered significant, **P<0.01, ***P<0.001, ****P<0.0001.

**Fig 10:**
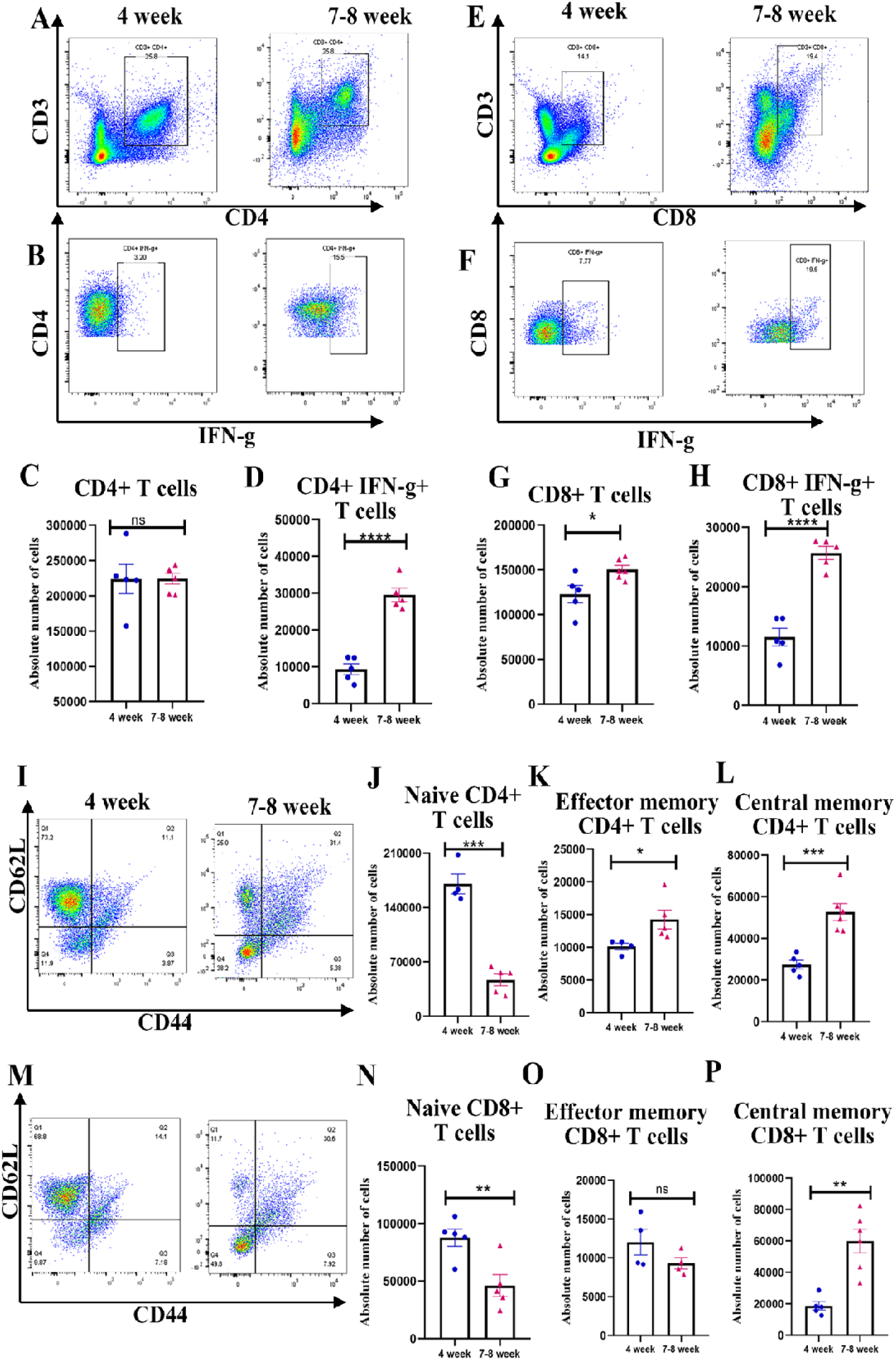
The priming activation and maturation of T cells were significantly enhanced in the young adult cohort at the innate-adaptive transition phase of neuroinflammation. On day 7 p.i., CLNs from MHV-RSA59-infected 4-week and 7-8-week-old C57BL/6 mice were harvested for flow cytometry analysis and stained for CD3, CD4, CD8, IFN-γ, CD62L, and CD44. Primary gating was performed on live cells, followed by singlets. Representative flow cytometry dot plots showing percentages of **A)** CD3+ CD4+ T cells, **B)** CD3+ CD4+ IFN-γ+ T cells. Graphical representation of the absolute cell numbers of **C)** CD4+ T cells, **D)** IFN-γ expressing CD4+ T cells. Representative flow cytometry dot plots showing percentages of **E)** CD3+ CD8+ T cells, **F)** CD3+ CD8+ IFN-γ+ T cells. Graphical representation of the absolute cell numbers of **G)** CD8+ T cells, **H)** IFN-γ expressing CD8+ T cells. **I)** Representative flow cytometry dot plots showing percentages of CD4+ CD62L+ CD44-(Naïve CD4+ T cells), CD4+ CD62L-CD44+ (Effector memory CD4+ T cells), CD4+ CD62L+ CD44+ (Central memory CD4+ T cells). Graphical representation of the absolute cell numbers of **J)** Naïve, **K)** effector memory, and **L)** central memory CD4+ T cells. **M)** Representative flow cytometry dot plots showing percentages of CD8+ CD62L+ CD44-(Naïve CD8+ T cells), CD8+ CD62L-CD44+ (Effector memory CD8+ T cells), CD8+ CD62L+ CD44+ (Central memory CD8+ T cells). Graphical representation of the absolute cell numbers of **N)** Naïve, **O)** effector memory, and **P)** central memory CD8+ T cells. Results were expressed as mean ± SEM. The shown flow cytometry study is representative of one independent experiment. (n=4-6 per age group) *Asterisk represents statistical significance calculated using unpaired Student’s t-test, p<0.05 was considered significant, **P<0.01, ***P<0.001, ****P<0.0001.

Phenotypic analysis of naïve vs. antigen-experienced states revealed that on day 5 p.i., the pool of naïve CD4+ T cells (Fig. 9I, J) was comparable between the two age groups, whereas naïve CD8+ T cells (Fig. 9M, N) were already significantly reduced in young adults. By day 7 p.i., both the naïve CD4+ and CD8+ T cell pools (Fig. 10I, J and Fig. 10M, N) were markedly lower in the young adult group than in the juvenile group. Crucially, effector CD4+ T cells (Fig. 9I, K and Fig. 10I, K) were significantly enriched in the CLNs of young adult mice on both days 5 and 7 p.i. Effector CD8+ T cells were significantly higher in young adults on day 5 p.i. (Fig. 9M, O) but became comparable between the groups by day 7 p.i. (Fig. 10M, O).

Additionally, central memory CD4+ (Fig. 9I, L and Fig. 10I, L) and CD8+ T cell populations (Fig. 9M, P and Fig. 10M, P) were significantly increased in young adult mice compared to juveniles across both timepoints. CLN T cell activation and early memory indicators were higher in young adults at days 5-7 p.i.The reduced pool of naïve T cells alongside the expansion of effector and central memory phenotypes implies highly efficient early activation and priming within adult secondary lymphoid organs, generating effector immune populations poised to migrate to the infected CNS.

### Regulatory T cells (Tregs) in the CLN and their accumulation within the CNS were significantly higher in young adult mice during early infection

To investigate the balancing counter-regulatory mechanisms, the kinetics of CD4+CD25+FOXP3+ regulatory T cells (Tregs) were examined. Accordingly, the quantitative profiling of Tregs revealed that their accumulation within the brain begins during the acute phase (day 5 p.i.) and peaks during the innate-adaptive transition phase at day 7 p.i. (Fig. 11A-D). Notably, the absolute numbers of Tregs were significantly higher in the brains of young adult mice compared to juveniles at both time points.

**Fig 11:**
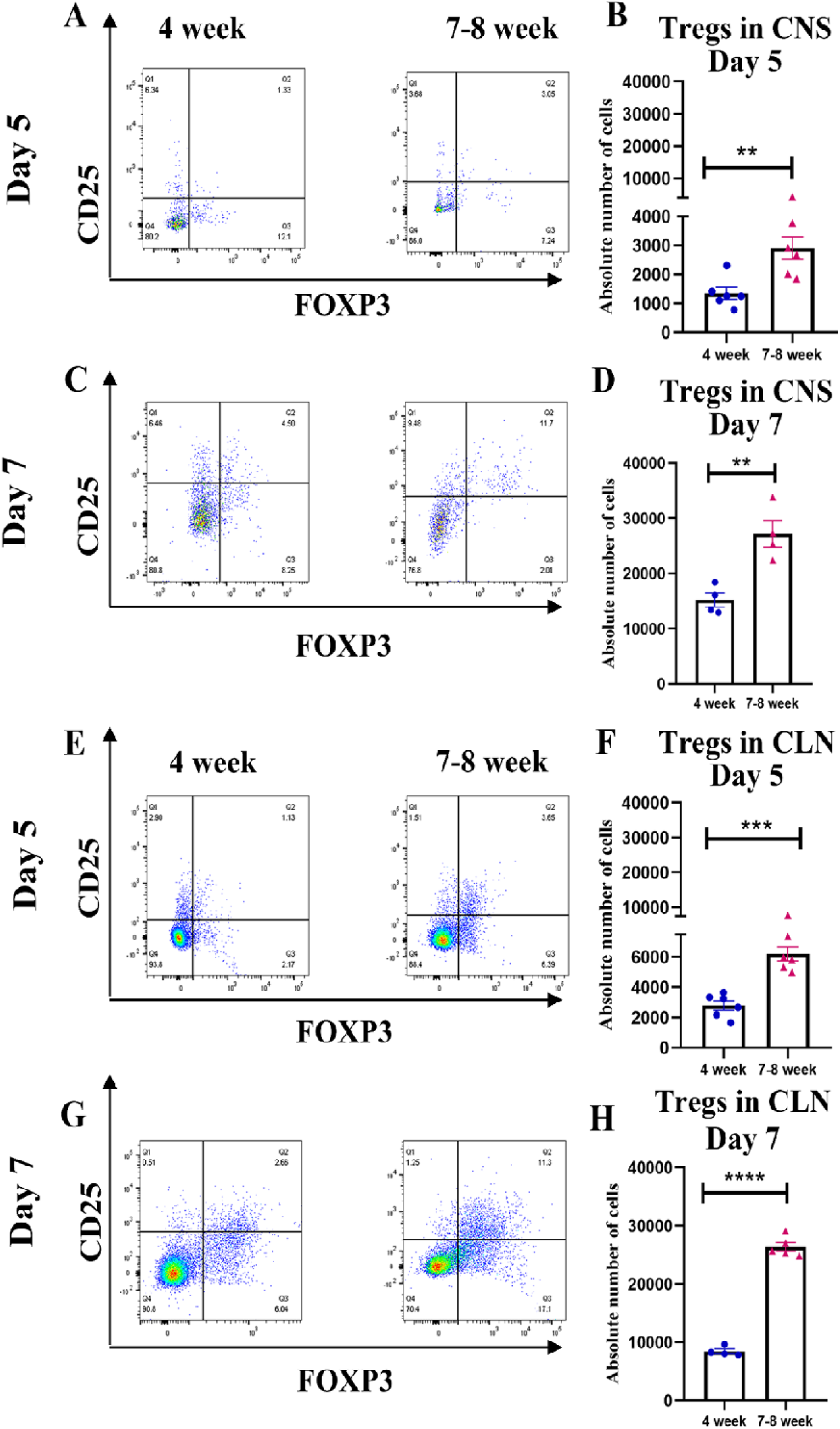
The accumulation of Tregs in the CNS, as well as its abundance in the CLN, was significantly higher in the young adult mice: On days 5 and 7 p.i., brains from MHV-RSA59-infected 4-week and 7-8-week-old C57BL/6 mice were harvested for flow cytometry analysis and stained for CD45, CD3, CD4, CD25, and FOXP3. CLNs were also harvested for flow cytometry analysis and stained for the previously mentioned markers except CD45. Primary gating was performed on live cells, followed by singlets. **A, C)** Representative flow cytometry dot plots showing percentages of CD45 hi CD3+ CD4+ CD25+ FOXP3+ T cells (Tregs) in CNS. **B, D)** Graphical representation of the absolute cell numbers of infiltrating Tregs. **E, G)** Representative flow cytometry dot plots showing percentages of CD3+ CD4+ CD25+ FOXP3+ T cells (Tregs) in CLN. **F, H)** Graphical representation of the absolute cell numbers of Tregs in CLN. Results were expressed as mean ± SEM. The shown flow cytometry study is representative of one independent experiment. (n=4-6 per age group) *Asterisk represents statistical significance calculated using unpaired Student’s t-test, P<0.05 was considered significant, **P<0.01, ***P<0.001, ****P<0.0001.

To determine whether this CNS accumulation reflected age-dependent differences in peripheral generation or expansion, Tregs within the draining CLNs were analyzed in parallel. Absolute cell counts confirmed that Treg populations were significantly larger in the CLNs of young adult mice on both days 5 and 7 p.i. compared to their juvenile counterparts (Fig. 11E-H), with Tregs generation increasing significantly in the CLN by day 7 p.i.

### Pro-inflammatory and anti-inflammatory cytokine profiles and the antiviral effector Ifit2 were significantly upregulated in the brains of young adult mice

To map the cytokine environment driving these cellular shifts, whole-brain transcript expression was analyzed via qRT-PCR. The results demonstrated that the expression of pro-inflammatory cytokines (TNF-α, IL-6) and key antiviral interferons (IFN-α, IFN-β, and IFN-γ) was significantly higher in young adult mice at the acute-innate stage (day 5 p.i.) compared to juveniles (Fig. 12A-E). Concurrently, the anti-inflammatory cytokine TGF-β (Fig. 12F) was significantly upregulated in the young adult CNS, while IL-10 transcripts (Fig. 12G) remained statistically equivalent between the two age groups. Furthermore, transcriptional analysis of critical leukocyte-recruiting chemokines (CXCL1, CXCL9, CXCL10, and CCL5) revealed significantly higher expression across all chemokines examined (Fig. 12H-K) in the young adult cohort.

**Fig 12:**
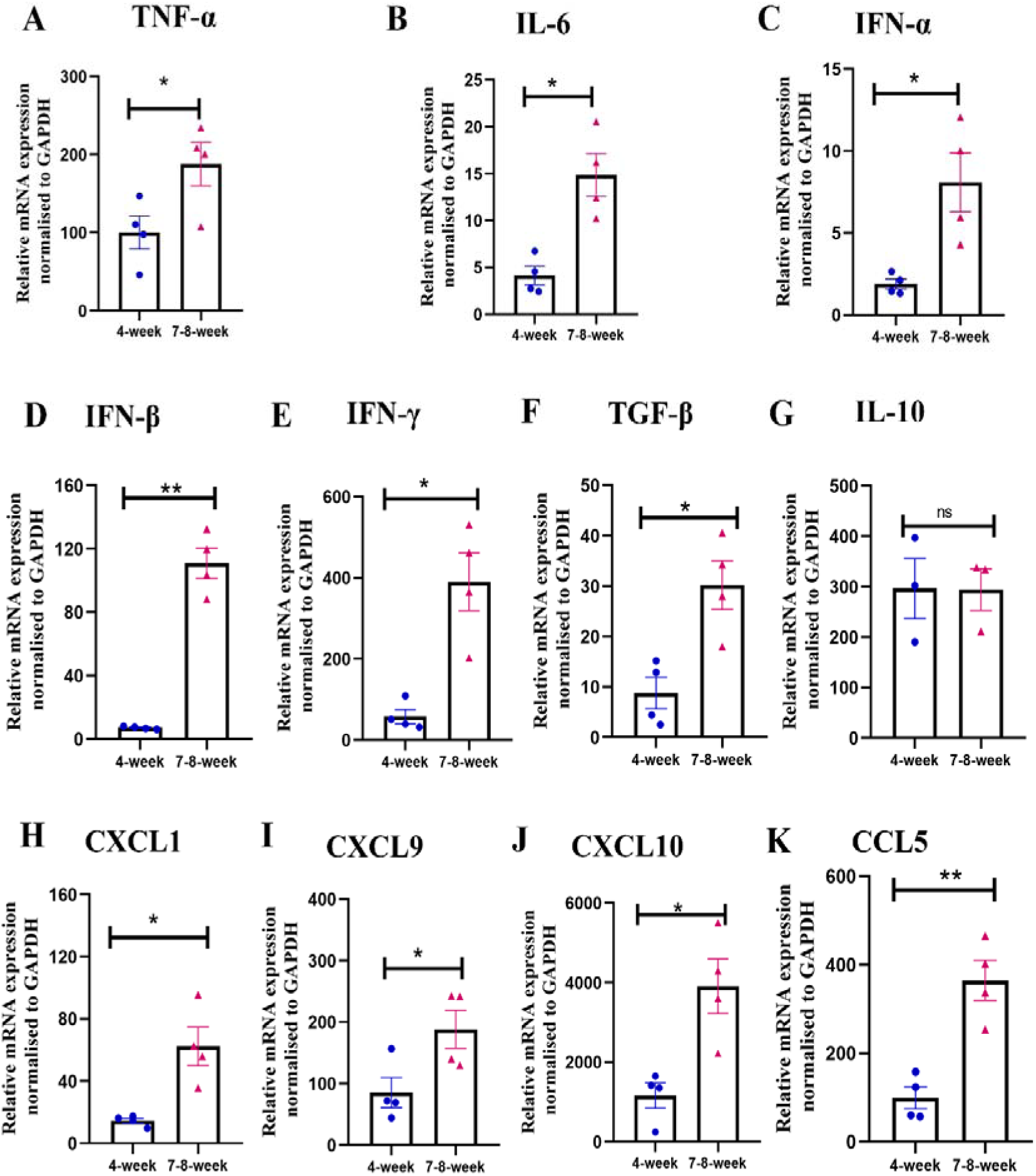
The young adult mice showed significantly higher expression of cytokines and chemokines in the CNS: (A-K) On day 5 p.i., the abundance of TNF-α, IL-6, IFN-α, IFN-β, IFN-γ, TGF-β, IL-10, CXCL1, CXCL9, CXCL10, CCL5 at transcript level was determined in brain lysates from MHV-RSA59-infected 4-week and 7-8-week C57BL/6 mice by qRT-PCR. The mRNA levels were normalized to *Gapdh*, which was used as the housekeeping control. The fold changes were compared to the mock-infected control group and expressed as mean±SEM. The shown qRT-PCR study is representative of one independent experiment. (n=3-4 per age group). *Asterisk represents statistical significance calculated using unpaired Student’s t-test, P<0.05 was considered significant, **P<0.01.

Because type I interferons (IFN-α and IFN-β) were highly upregulated in the young adult CNS (Fig. 12C, D), the expression of the downstream interferon-stimulated gene (ISG) *Ifit2* (IFN-induced protein with tetratricopeptide repeats 2) was also examined. Longitudinal qRT-PCR analysis of *Ifit2* transcripts revealed significantly higher expression levels on days 3, 5, and 7 p.i. in the young adult mouse group compared to juveniles. By day 30 p.i., when replicating viral particles are cleared from the system, *Ifit2* transcripts are expressed only at a basal level in mice from both age groups (Fig. 13A). Complementary immunoblotting demonstrated a significantly higher expression of Ifit2 protein on day 5 p.i. in young adult mice, whereas by day 7 p.i., protein levels became equivalent between the two groups (Fig. 13B-F). The temporal correlation between elevated Ifit2 expression during the acute phase and reduced infectious viral titers at day 7 p.i. in young adult mice is consistent with Ifit2 contributing to viral restriction, although direct testing in Ifit2-deficient backgrounds will be required to confirm this important antiviral role.

**Fig 13:**
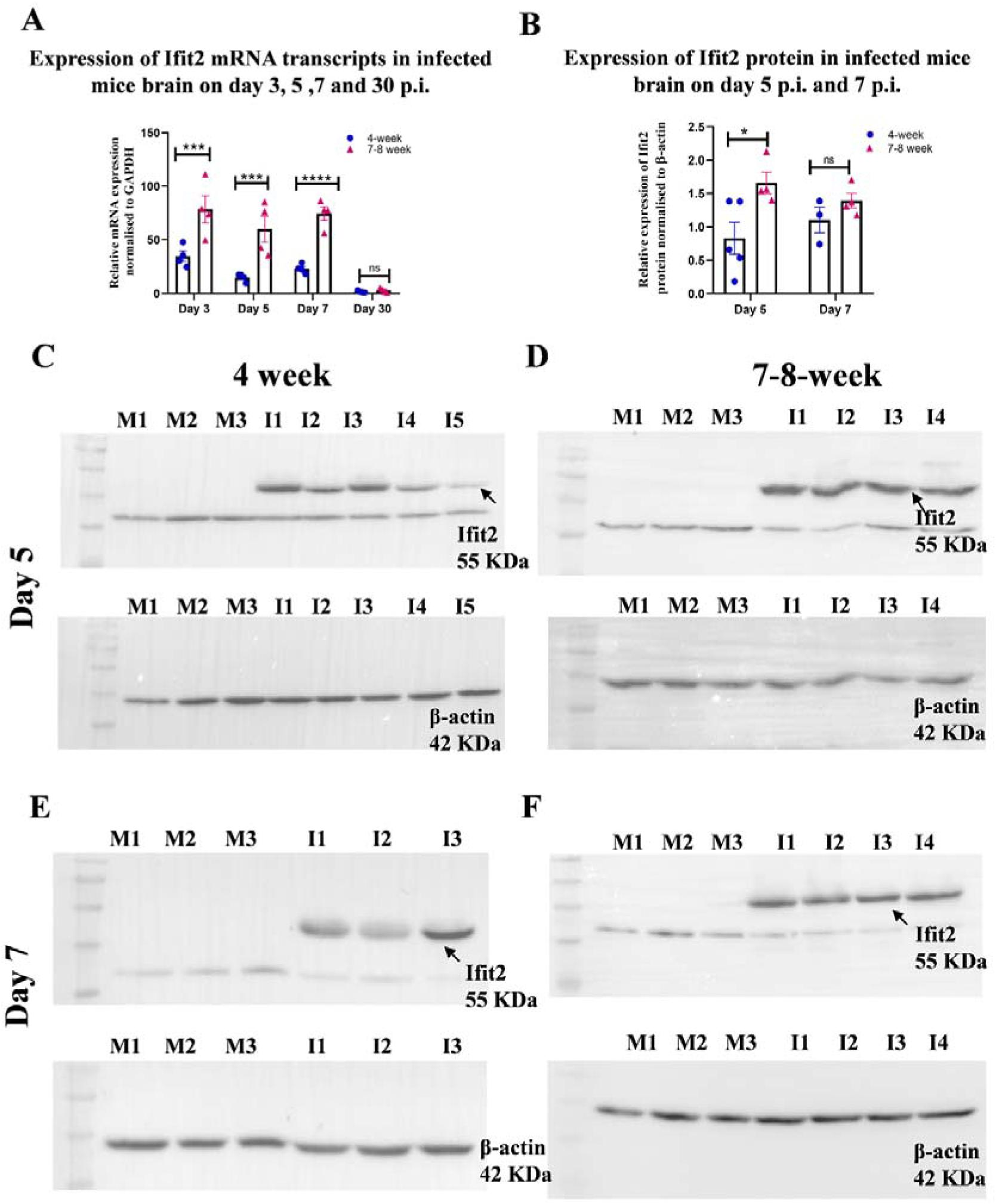
The higher expression of Ifit2 in the young adult mice at the innate acute phase restricts viral replication and causes increased viral clearance. **A)** Abundance of *Ifit2* transcript was determined in brain lysates from MHV-RSA59-infected 4-week and 7-8-week C57BL/6 mice on days 3, 5, 7, and 30 p.i. by qRT-PCR. The *Ifit2* mRNA levels were normalized to *Gapdh*, which was used as the housekeeping control. The fold changes were compared to the mock-infected control group. **B)** Histogram shows the relative expression of Ifit2 normalized to β-actin. The fold changes were compared to the mock-infected control group. **(C-F)** Representative immunoblots showing Ifit2 (55 KDa), indicated by an arrow, and β-actin (42 KDa) expression in mice brain homogenates collected on day 5 p.i. and day 7 p.i. respectively. Results were expressed as mean ± SEM. (n=3-5 per age group). *Asterisk represents statistical significance calculated using an unpaired Student’s t-test; P<0.05 was considered significant. ***P<0.001, ****P<0.0001.

In summary, our findings demonstrate that young adult mice exhibit a significantly higher resistance to MHV-RSA59-induced chronic demyelination compared to juvenile mice. This protective phenotype is driven by robust, early innate and adaptive immune responses characterized by heightened glial cell activation, increased myeloid and lymphoid cell infiltration, elevated pro-inflammatory cytokine and interferon production, and enhanced interferon-stimulated gene (ISG) expression. Collectively, these coordinated immune mechanisms achieve superior local viral control by day 5-7 p.i., effectively preventing the early-onset demyelination pathologically observed in juvenile cohorts as early as day 5 post-infection.

## Discussion

In the post-COVID-19 era, the pandemic has further amplified awareness of coronavirus neurological diseases(16, 17, 27, 39, 40) and the emergence of multiple SARS-CoV-2 variants has been associated with distinct differences in virulence, transmissibility, and immune response profiles across various age groups(41–44). These clinical findings prompted us to hypothesize that age-dependent immune maturation may be a critical factor in protecting the host from β-coronavirus infectivity and mitigating the severity of neuropathology, specifically, demyelination. To test this hypothesis, this study attempted to investigate the age-dependent immune maturation and its implications for murine coronavirus pathogenesis during both the acute and chronic phases of infection. MHV and SARS-CoV-2 belong to the same family of β-coronaviruses, and many of the immune evasion strategies and neuropathogenic mechanisms described in MHV models have direct counterparts in human coronavirus infections; hence, experimental insights into murine-coronavirus-induced neuroinflammation and demyelination are crucial for understanding the potential neurological consequences observed in SARS-CoV-2 patients(45).

This study used juvenile (4-week-old) and young adult (7-8-week-old) mouse cohorts to investigate the neurological sequelae of coronavirus infection and examine how the immune response varies in an age-dependent manner following murine β-coronavirus infection, specifically assessing disease severity, neuroinflammation resolution, immune activation, and long-term sequelae such as demyelination. Demyelination analysis data revealed a marked attenuation of demyelination pathology in young adult mice at the chronic stage (day 30 p.i.) compared to juvenile counterparts. Reduced demyelination was associated with a significant decrease in the prevalence of activated microglia and macrophages in the spinal cord white matter. Given the established role of these immune cells in actively stripping myelin (33) we propose that the heightened susceptibility in juveniles may stem from a dysregulated immune response. Such dysregulation could prevent microglia/macrophages from returning to homeostasis, thereby perpetuating their activated phenotype and culminating in persistent myelin damage. The chronic inflammatory demyelination pathology observed only in juvenile mice provided insights to pursue further mechanistic studies and prompted us to investigate disease progression at the initial disease stages, when infectious virus particles are prevalent in the system.

To understand the pathophysiological mechanisms underlying the enhanced demyelination observed in juvenile mice at the chronic stage, disease pathology during the early stages of infection was subsequently investigated. The weight loss and disease severity results revealed a comparable weight reduction in both age groups, while the clinical scores revealed a significantly reduced disease severity in young adult mice at the innate-adaptive transition stage. The low disease severity was attributed to significantly lower viral loads in young adult mice by the innate-adaptive transition stage, implying that this age group was better suited to combat the virus infection. Hence, it was imperative to investigate neuroinflammation and CNS glial cell activation on MHV-RSA59 infection. Encephalitis, marked by perivascular cuffing and microglial nodules formation and the activation of microglia and astrocytes, was significantly enhanced in young adult mice compared to juveniles on days 5 and 7 p.i., resulting in an increased neuroinflammation, an outcome of a protective immune response exhibited by glial cells directed towards reducing viral replication and spread.

In mock-infected juvenile and young adult mice, the numbers and activation of brain-resident microglia and peripheral infiltrating leukocytes were comparable between the two age groups. Upon infection, microglia and astrocytes get activated and secrete different cytokines and chemokines(46–49). During the acute phase, elevated levels of TNF-α and IL-6 in young adult mice trigger inflammatory cascades and probably result in significantly greater neutrophil and monocyte/macrophage infiltration compared to juvenile mice. Additionally, elevated CXCL1 production in young adult mouse brains could also drive increased neutrophil migration(50, 51). The increased numbers and heightened activation of CX3CR1+ and MHC II+ brain-resident microglia in young adult mice indicated an amplified pro-inflammatory response that is characteristic of effective antiviral host immunity.

In virus-induced neuroinflammation, CX3CR1 signaling plays a complex role and can be protective or detrimental(52, 53). Consistent with previous reports demonstrating a protective role for CX3CR1-activated microglia in promoting lymphocyte infiltration into the CNS(37), our recent findings suggest that increased CX3CR1 expression in young adult mice reflects an age-dependent augmentation of the host-protective immune response, leading to enhanced activation and recruitment of monocytes/macrophages and activated T cells, including CD4+ T cells, NKT cells, and IFN-γ-expressing CD4+ and CD8+ T cells. Upon entering the CNS, monocytes and macrophages show heightened MHC II expression, facilitating reactivation of virus-specific CD4+ and CD8+ T cells and resulting in IFN-γ production(54).

The generation of IFN-γ-producing CD4+ and CD8+ T cells was also significantly higher in the CLNs of young adult mice. The mechanism underlying the greater infiltration of IFN-γ-producing CD4+ and CD8+T cells into the CNS of young adult mice may involve an IFN-γ-driven amplification loop. The virus-specific IFN-γ+ T cells, upon entering the CNS, reactivate CNS-resident glial cells and might stimulate them to produce CXCL9/10(54–57). The production of CXCL9 and CXCL10 is significantly higher in the CNS of young adult mice compared to juvenile mice. The chemokine gradient promotes further recruitment of CXCR3+ effector T cells into the CNS, thereby amplifying the immune response. In young adult mice, this positive feedback loop may be established efficiently, leading to sustained effector T cell accumulation and enhanced viral control, a dynamic that may be compromised in the juveniles.

The ISG Ifit2 is known to exert antiviral activity against several viruses, including West Nile virus, Sendai virus, vesicular stomatitis virus (VSV), and mouse hepatitis virus (MHV)(58). In the context of MHV-RSA59 infection, previous studies have shown that Ifit2 confers antiviral protection during both the acute and chronic stages of infection. Ifit2 ablation impairs CX3CR1 expression in monocyte/microglia/macrophages, leading to reduced T cell trafficking in the CNS and, ultimately, impaired viral clearance(37, 38). Taking insights from previous studies, we sought to determine whether the expression of Ifit2 varied in an age-dependent manner. The increased expression of type I interferons, such as IFN-α and IFN-β, in young adult mice, and the concomitant upregulation of the ISG Ifit2, was one of the factors associated with a pronounced reduction in viral load, indicating an effective antiviral state. This observation is consistent with emerging evidence that microglia are initial inducers of IFN-α/β, which acts in both an autocrine and paracrine fashion to broadly upregulate an antiviral gene expression program(59).

Immunophenotyping in the CLN on days 5 and 7 p.i. revealed that the effective resolution of viral infection in young adult mice is linked to distinct T cell dynamics. The significant decrease in naïve CD4+ and CD8+ T cells in young adult mice relative to juveniles was accompanied by a significant expansion of effector and central memory T cell populations. The increased central memory T cell response indicates the development of a robust, effective memory response ensuring long-term protection. The persistence of significantly high numbers of naïve T cells and lower numbers of effector and central memory T cells in juvenile mice may reflect a delayed or suboptimal T cell-mediated immune response, consistent with the concept that suboptimal antigen-specific immunity underlies age-related susceptibility to severe disease outcomes.

Neuroinflammation is a double-edged sword that becomes destructive when immune cells get overactivated. Consistent with previous reports showing that Tregs play a protective role in MHV infection by limiting Th1 immunopathology(60), our findings show that increased Treg production in the CLN and enhanced CNS infiltration are critical for restoring homeostasis. Tregs release anti-inflammatory cytokines, such as IL-10 and TGF-β,(61–63) with significantly higher TGF-β levels in young adult mice. In a previous study, virus-specific Tregs have been shown to inhibit effector T cell proliferation and migration from the draining lymph node, diminish microglia activation, and decrease the number and function of effector T cells in the infected brain, thereby reducing mortality and morbidity without affecting virus clearance(60). In another study, virus-specific Tregs have been shown to persist long-term after murine infection, maintaining FOXP3 expression for at least 180 days post-infection, suggesting the establishment of a durable memory Treg compartment(64).

Although both effector T cell infiltration and Tregs accumulation are high in young adult mice at the innate-adaptive transition stage, we propose that Tregs may persist even at later stages of infection and serve as a critical regulator. The continued Treg infiltration after viral burden declines likely prevents excessive effector T cell activity, thereby preventing immune overactivation and progressive tissue damage. Previous studies have shown that CD4+ T cells exert protective functions in MHV-RSA59 infection by modulating microglial/macrophage activation, thereby reinstating homeostasis in the CNS(36, 65). The reduced demyelination observed in young adult mice at the chronic stage may be attributable to the enhanced and timely accumulation of Tregs within the CNS, which prevents microglia/macrophage overactivation(66). By releasing anti-inflammatory cytokines such as TGF-β, Tregs may expedite the resolution of neuroinflammation by promoting a phenotypic shift in microglia and macrophages from a pro-inflammatory to a reparative state. This reparative shift is supported by significantly higher CD206 expression in the spinal cords of young adult mice and lower expression of microglial phagocytic markers such as TREM-2 and CD68, consistent with a transition away from the destructive, myelin-stripping phenotype.

Taken together, our data support an integrative model in which the maturity of the host immune system at the time of coronavirus infection is a primary determinant of long-term neuropathological outcome. In young adult mice, a robust and temporally coordinated immune response characterized by efficient type I interferon and Ifit2 induction, increased microglia/macrophage CD40 activation, amplified CX3CR1-driven microglial activation, extensive recruitment of IFN-γ-producing T cells via the CXCL9/CXCL10-CXCR3 axis, rapid effector and memory T cell differentiation, and timely Treg-mediated immunoregulation collectively enables effective viral control and prevents exacerbated neuroinflammation. In juvenile mice, an impaired host immune response when challenged with neurotropic coronaviruses results in a failure to establish these protective immune circuits, culminating in extensive demyelinating plaque formation in the spinal cord white matter. In summary, reduced demyelination in young adult mice is due to optimal age-dependent immune maturation and a protective host immune response. In conclusion, these findings indicate that age is a crucial factor in imparting protective immunity against coronavirus infection and provide a vital first step toward dismantling the precise cellular and immunological mechanisms by which an older, more mature immune system restricts coronavirus-induced neurodegeneration.

## Material and Methods

### Virus, mice inoculation, and experimental design

The virus used in the study (MHV-RSA59) is a neurotropic demyelinating isogenic recombinant strain of MHV-A59(35, 67). Four-week-old juvenile and seven-eight-week-old young-adult wild-type (WT) C57BL/6 male mice inbred at IISER-Kolkata were used for the study. The mice were intracranially inoculated with the viral inoculum, 20,000 PFU (50% of LD50) of the MHV-RSA59 strain(67). Mock-infected control mice were inoculated with sterilized (PBS+0.75% BSA) in a final volume of 20μl. All experimental mice were daily monitored for the appearance of disease signs and symptoms. The clinical disease severity was graded in accordance to the following scale: 0-no disease symptoms, 0.5-ruffled fur, 1.0-hunched back position with mild ataxia and slightly slower movement, 1.5-hunched back position with mild ataxia, hindlimb weakness, restricted movement, 2-ataxia, balance problem and/or partial paralysis, still capable of moving, 2.5-complete paralysis in one leg, motility issue, movement with difficulties, 3-severe hunchback, paralysis in both hindlimbs and movement is severely compromised, 3.5-complete paralysis with no movement and moribund, 4-dead(36, 68). For histopathological and qRT-PCR experiments, mice were sacrificed at the innate-acute stage of infection (day 5 p.i.), innate-adaptive transition stage (day 7 p.i.), and chronic stage (day 30 p.i.). For immunophenotyping and immunoblotting experiments, mice were sacrificed on days 5 and 7 p.i. For viral titer assays, mice were sacrificed on days 3, 5, and 7 p.i.

### Estimation of viral titer as a readout of viral replication

Whole brain tissues (approximately 450 mg) were homogenized using Tenbroeck tissue homogenizers in 4 ml of RPMI containing 25 mM HEPES (pH 7.3). Centrifugation was performed at 450g for 10 min, and the cell pellets were used for flow cytometry, while the supernatant was separately stored at -80°C for viral plaque assay. MHV-RSA59 titers in the brain homogenate were determined by standard plaque assay protocol on monolayers of L2 cells using the formula: plaque-forming units (PFUs) = (no. of plaques X dilution factor/ml/gram of tissue) and expressed as log10 PFUs/gram of tissue(36).

### Histopathology and immunohistochemical analysis

In accordance with our lab group’s protocol(36), brain and spinal cord tissues were harvested following transcardial perfusion with 20 ml PBS. Tissues were fixed in 4% paraformaldehyde for 36-48 hours, after which they were processed through increasing concentrations of ethyl alcohol, xylene, and paraffin wax. Tissues were embedded in paraffin, and 5 μm transverse sections (spinal cord) and mid-sagittal sections (brain) were obtained for staining with Hematoxylin and Eosin for histopathological analysis. Moreover, spinal cords were also stained with Luxol fast blue (LFB) to determine the extent of demyelination(33, 68). Immunohistochemical staining of the brain and spinal cord tissue sections was performed using the avidin-biotin immunoperoxidase technique as described previously (Vector Laboratories) using 3,3’-Diaminobenzidine as the substrate (catalog# SK-4100, PK-4001). The following primary antibodies were used: 1:600 dilution of anti-Iba1 (Wako, catalog#019-19741), 1:600 dilution of anti-GFAP (Sigma-Aldrich, MO, USA, catalog# G4546). Control slides from mock-infected mice were stained in parallel.

### Quantification of histopathological sections

Image analysis was conducted using the densitometric thresholding tool in Fiji (ImageJ, NIH Image, Scion Image). For Iba1 and GFAP immunohistochemical staining, images were acquired at defined magnifications (4X for brain sections and 10X for spinal cord sections) to ensure visualization of the entire tissue section within a single frame. RGB images were deconvoluted into three color channels to isolate and subtract DAB-specific staining from the background hematoxylin signal. The boundaries of each brain and spinal cord section were digitally delineated, and the total tissue area was calculated in μm². A threshold value was set for each image to ensure that all antibody-marked cells were included. The amount of Iba1 and GFAP staining was termed as “% area of staining.” (36, 65, 69)

The % area of demyelination was determined in LFB-stained spinal cord sections from each mouse and analyzed using Fiji (ImageJ, NIH Image, Scion Image)(69). The white matter regions in each spinal cord cross-section were marked and calculated by adding together the dorsal, ventral, and anterior white matter areas in each section. Also, the total area of demyelinated regions was outlined and compiled for each section. The percentage area of demyelination per spinal cord section per mouse was calculated (2-3 sections from the cervical, thoracic, and lumbar regions of the spinal cord/mouse were examined).

### Protein isolation and immunoblot analysis

In accordance with our lab group’s protocol(70), approximately 150mg of brain tissue was collected following transcardial perfusion with PBS and flash-frozen in liquid nitrogen. Tissue was then lysed in 1 ml of RIPA buffer (0.1% SDS and 0.1% Triton X-100) with 1X complete mini protease inhibitor cocktail tablets (Roche, catalog# 11836153001) and phosphatase inhibitor cocktail: sodium orthovanadate (10 mM), sodium fluoride (10 mM), and sodium pyrophosphate (10 mM). Brain tissues were homogenized by trituration and sonication, and tissue lysate was centrifuged at 13,500 RPM for 30 min at 4°C, and the supernatant was collected as whole protein extract. Protein was quantified using Pierce BCA protein assay kit (Thermo Scientific, Rockford, IL, USA, catalog# 23225). Equal amounts of protein were loaded and resolved on SDS-PAGE, followed by transfer to polyvinylidene difluoride membranes (Millipore, Bedford, MA, catalog# IPVH00010) using transfer buffer (25mM Tris, 192mM glycine, and 20% methanol). The membrane was then blocked with 5% non-fat skimmed milk in TBST (Tris-buffered saline containing 0.1% v/v Tween-20) for 1 hour at room temperature, followed by incubation in primary anti-Ifit2 antibody (Thermo Scientific, 1:2500, catalog# PA3-845) and anti-β-actin antibody (Invitrogen, 1:10000, catalog# MA1-140) in blocking solution overnight at 4°C. Membranes were then washed three times in TBST, followed by incubation with HRP-conjugated secondary IgG. Membranes were subjected to washes in TBST, and immunoreactive bands were visualized using SuperSignal^TM^ West Pico PLUS Chemiluminescent Substrate (Thermo Fisher Scientific, catalog# 34578). Densitometric analyses of non-saturated membranes were carried out using a Syngene G:BOX chemidoc system and Image J software.

### Gene expression: RNA isolation, reverse transcription, and quantitative polymerase chain reaction

In accordance with our lab group’s protocol(70), RNA was extracted from brain or spinal cord tissues (flashfrozen) of MHV-RSA59 and mock-infected mice using the TRIzol isolation protocol following transcardial perfusion with PBS. The total RNA concentration was measured using a NanoDrop ND-2000 spectrophotometer. 1 µg of RNA was used to prepare cDNA using a High-Capacity cDNA Reverse Transcription Kit (Applied Biosystems, catalog# 4368814). Quantitative Real-time PCR analysis was performed using iTaq Universal SYBR Green Supermix qPCR kit (BioRad, catalog# 1725124) in a BioRad CFX Real-time PCR system (BioRad), maintaining the following conditions: initial denaturation at 95°C for 7 min, 40 cycles of 95°C for 10 sec, 60°C for 30 sec, and melting curve analysis at 60°C for 30 sec. Reactions were performed in quadruplets. Relative quantitation was performed using the comparative threshold (ΔΔCt) method. mRNA expression levels of target genes in MHV-RSA59 and mock-infected mice were normalized with Gapdh and expressed as relative fold change compared with their respective mock-infected controls. Primer sequences are listed in Table 1.

**Table 1:**
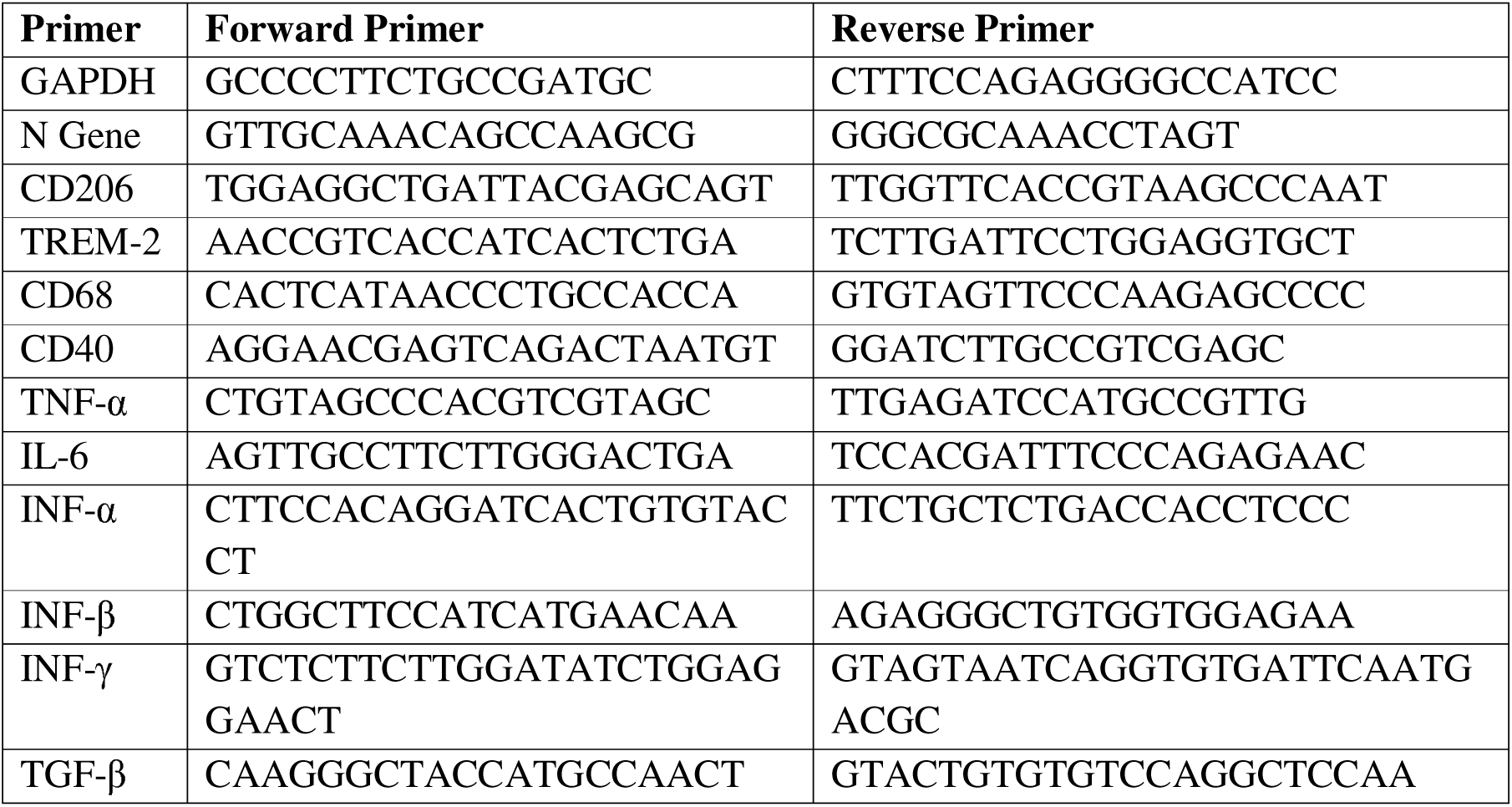
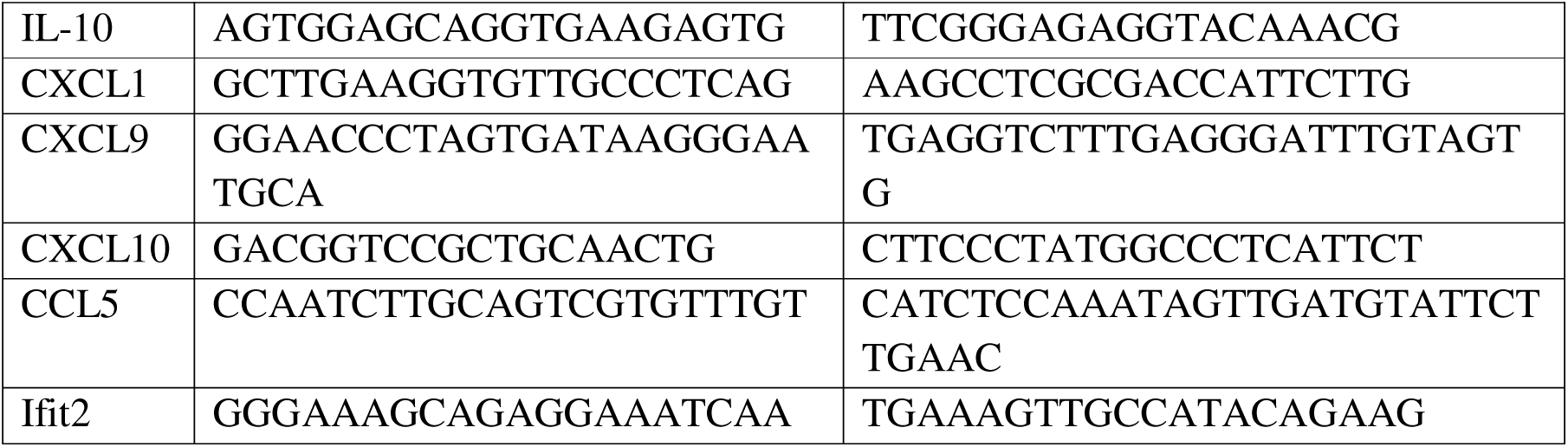
Sequence of primers used in the study.

### Flow cytometry analysis

In accordance with our lab group’s protocol(36, 37), mice were perfused transcardially with PBS, and whole brains were homogenized in 4 ml of RPMI containing 25 mM HEPES (pH 7.3), using Tenbroeck tissue homogenizers. Following centrifugation at 450g for 10 min, the cell pellets were resuspended in RPMI containing 25 mM HEPES, adjusted to 30% Percoll (Sigma), and underlaid with 1 ml of 70% Percoll. Following centrifugation at 850 g for 45 min at 4°C, cells were recovered from the 30%-70% interface, washed with RPMI, and suspended in FACS buffer (0.5% bovine serum albumin in Dulbecco’s PBS). Also, CLNs were harvested, homogenized in 5 ml RPMI containing 25 mM HEPES (pH 7.3), and passed through 70μm filters followed by 30μm filters to obtain single-cell suspensions. Following centrifugation at 450g for 10 min, cell pellets were resuspended in FACS buffer. Cells were counted using an automated cell counter (Invitrogen) to obtain the total number of leukocytes. Specific cell types in the brain and CLN were identified by staining with fluorochromes like fluorescein isothiocyanate (FITC), phycoerythrin [(PE), (PECy7)], peridinin chlorophyll protein [(PerCpCy5.5)], allophycocyanin [(APC), (APCCy7)] and violet excitable dyes [(V450), (V500), (BV421)] conjugated mAb for 45 min at 4°C in FACS buffer. Expression of surface and intracellular markers was characterized with mAbs conjugated with specific fluorochromes as mentioned in Table 2. Intra-cellular staining was performed for FOXP3 after fixation and permeabilization (BD catalog# 562574). Intra-cellular staining was performed for IFN-γ after fixation and permeabilization (BDcatalog# 554714). Samples acquisition was performed on a BD FACSVerse flow cytometer and BD FACSAria III Flow Cytometer (BD Biosciences) and analyzed on FlowJo 10 software (Treestar, Inc., Ashland, OR).

**Table 2:** Antibodies used for flow cytometry experiments.

| Marker | Clone | Brand |
| --- | --- | --- |
| CD45 | Ly-5 | BD Biosciences |
| CD11b | M1/70 | BD Biosciences |
| CX3CR1 | SA011F11 | BioLegend |
| MHC II | 2G9 | BD Biosciences |
| Ly6G | 1A8 | BD Biosciences |
| CD3 | 145-2C11 | BD Biosciences |
| CD4 | GK1.5 | BD Biosciences |
| CD8 | 53-6.7 | BD Biosciences |
| NK1.1 | PK136 | BD Biosciences |
| CD44 | IM7 | BD Biosciences |
| CD62L | MEL-14 | BD Biosciences |
| CXCR3 | 1C6/CXCR3 | BD Biosciences |
| CD25 | PC61 | BD Biosciences |
| FOXP3 | G155-178 | BD Biosciences |
| IFN-γ | XMG1.2 | BD Biosciences |

Gating strategy in brain: Initially, the live cell population was separated from debris using the forward scatter area (FSC-A) and side scatter area (SSC-A). Subsequently, singlets were selected based on FSC-A and forward scatter height/width (FSC-H or FSC-W). This initial selection process was applied uniformly across all experiments. Gating on CD45, a pan-leukocyte marker, allowed for the distinction between (CD45-High) CD45hi peripheral immune cells and (CD45-low) CD45lo brain-resident immune cells. Myeloid cells were identified by CD11b expression. CD11b gating was applied to CD45lo cells to specify microglia. Among the peripheral leukocytes, neutrophils and monocyte/macrophages were distinguished within the CD45hi population through concurrent gating of CD11b and Ly6G. Both neutrophils and macrophages express CD11b, but Ly6G is only expressed by neutrophils. The lymphocyte population was isolated from the CD45hi population by applying a concurrent CD3 (pan-T cell marker) and CD4 gating to distinguish CD4+ T cells, while CD3 and CD8 gating were applied after CD45hi to distinguish CD8+ T cells. Further, the expression of activation markers was checked on myeloid antigen-presenting cells. CX3CR1+ and MHCII+ cells were gated separately from microglia (CD45lo CD11b+ CX3CR1+/MHC-II+) and infiltrating monocyte/macrophages (CD45hi CD11b+ Ly6G-CX3CR1+/MHC-II+). Effector CD4+ and CD8+ T cells were distinguished by an additional gating of CXCR3 on CD4+/CD8+ T cells. IFN-γ-expressing CD4+ and CD8+ T cells were distinguished by an additional gating of IFN-γ on CD4+/CD8+ T cells. Regulatory T cells (Tregs) were distinguished by an additional concurrent gating of CD25 and FOXP3 on CD4+ T cells. NKT cells were distinguished from the CD45hi population by applying a concurrent CD3 and NK1.1 gating.

Gating strategy for CLN: Similar to the brain, the live cell population was separated from debris using the forward scatter area (FSC-A) and side scatter area (SSC-A). Subsequently, singlets were selected based on FSC-A and forward scatter height/width (FSC-H or FSC-W). CD4+ and CD8+ T cells were gated from singlets by concurrent gating on CD3 and CD4/CD8. Finally, memory populations of CD4+ and CD8+ T cells in the CLN were analyzed by a concurrent gating of CD44 (effector cell marker) and CD62L (homing marker).

### Statistical analysis

Values were represented as mean ± standard errors of the mean (SEM). Values were subjected to unpaired Student’s t-tests with Welch’s correction or two-way ANOVA with multiple comparison tests (Sidak’s multiple comparison test) for calculating the significance of differences between the means. All statistical analyses were performed using GraphPad Prism 8 software (La Jolla, CA). A P-value of < 0.05 was considered statistically significant.

## Acknowledgements

We thank the State-of-the-Art Animal Facility, Indian Institute of Science Education and Research Kolkata (IISER-Kolkata), for providing all mice used in this study. We also thank the Department of Biological Sciences, IISER-Kolkata, for access to the flow cytometry facility, and Mr. Tamal Ghosh for his assistance with flow cytometry data acquisition. We thank Ms. Sreetama Bhaduri for her contribution to sample preparation for flow cytometry experiments. We acknowledge IISER-Kolkata for institutional support. Fellowship support for S.G. and B.H. was provided by the University Grants Commission (UGC), India.

## Ethics Statement

All animal experiments were conducted in accordance with the guidelines established by the Committee for the Control and Supervision of Experiments on Animals (CCSEA), India. The experimental protocol was reviewed and approved by the Institutional Animal Ethics Committee (IAEC) of the Indian Institute of Science Education and Research Kolkata (IISER-Kolkata), Protocol no-IISERK/IAEC/AP/2019/45, IISERK/IAEC/AP/2024/125, IISERK/IAEC/AP/2026/181. All procedures for animal experiments were performed in full compliance with institutional animal ethics. Every effort was made to minimize pain, distress, and discomfort through appropriate handling techniques, environmental enrichment, and adherence to humane endpoints. The number of animals used was limited to the minimum required to achieve statistical significance, in strict accordance with the 3Rs principles of Replacement, Reduction, and Refinement in biomedical research.

## Author Contributions

Satavisha Ghosh (S.G); Conceptualization, Data curation, Formal analysis, Investigation, Methodology, Software, Validation, Visualization, Writing-original draft, Writing-review and editing. Bishal Hazra (B.H); Data curation, Methodology, Software, Visualization. Subhajit Das Sarma (S.D.S); Methodology. Debanjana Chakravarty (D.C); Review and editing. Jayasri Das Sarma (J.D.S); Conceptualization, Formal analysis, Funding acquisition, Investigation, Project administration, Resources, Supervision, Validation, Visualization, Writing-original draft, Writing-review and editing.

## Data availability

The data supporting this study are available and can be obtained from the corresponding author.

---

**Fig S1:**
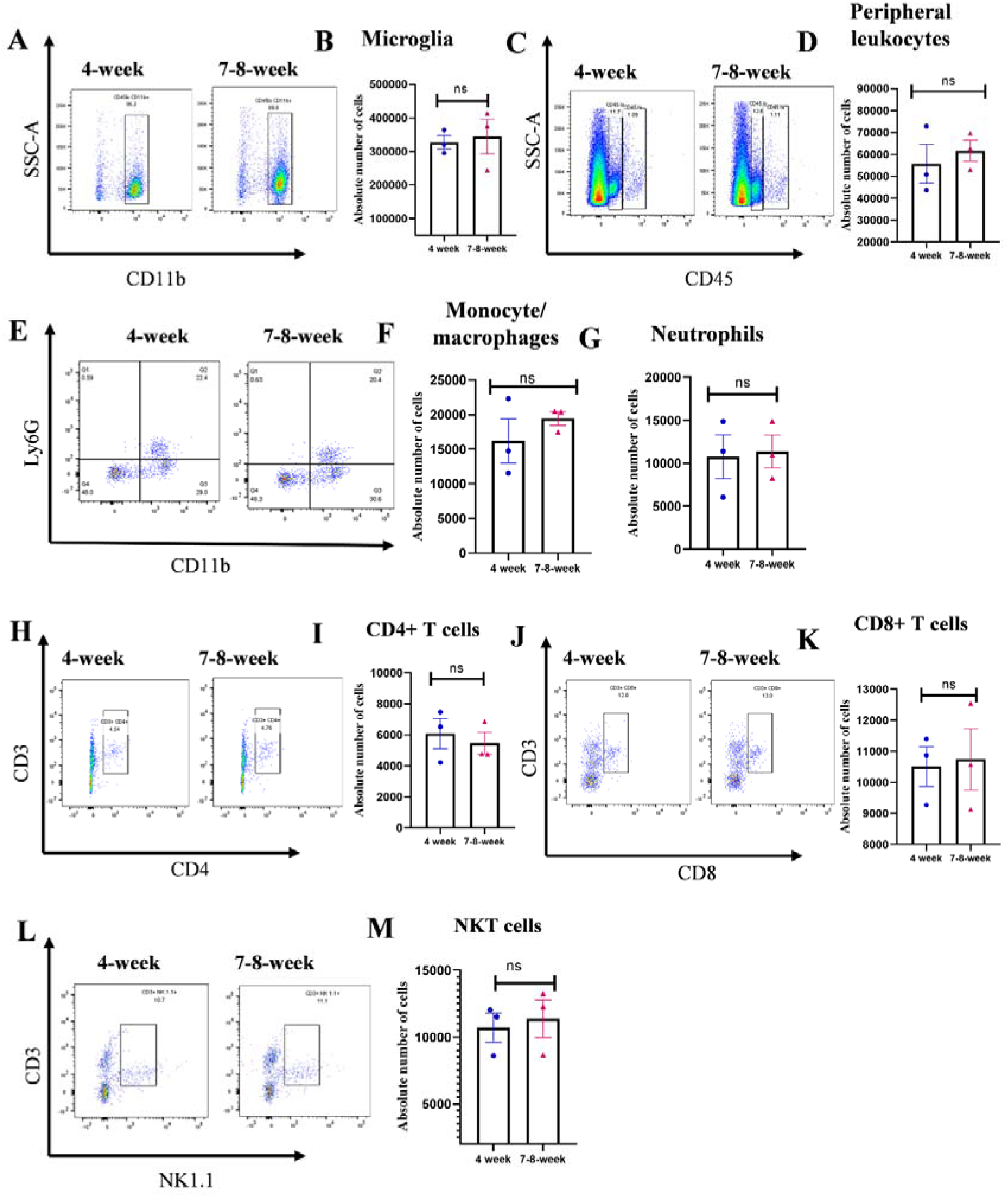
The number of brain-resident microglia and the infiltration of different peripheral leukocytes were comparable and equivalent in mock-infected juvenile and young adult mice groups. On day 5 p.i., brains from mock-infected 4-week and 7-8-week-old C57BL/6 mice were harvested for flow cytometry analysis and stained for CD45, CD11b, Ly6G, CD3, CD4, CD8, NK1.1. Primary gating was performed on live cells, followed by singlets. Representative dot plots and graphical representation of the absolute cell numbers of A, B) brain-resident microglia, C, D) peripheral leukocytes, E, F) monocyte/macrophages, E, **G)** neutrophils, **H, I)** CD4+ T cells, **J, K)** CD8+ T cells, **L, M)** NKT cells. Results were expressed as mean ± SEM. The shown flow cytometry study is representative of one independent experiment. (n=3 per age group) *Asterisk represents statistical significance calculated using an unpaired Student’s t-test; P<0.05 was considered significant.

**Fig S2:**
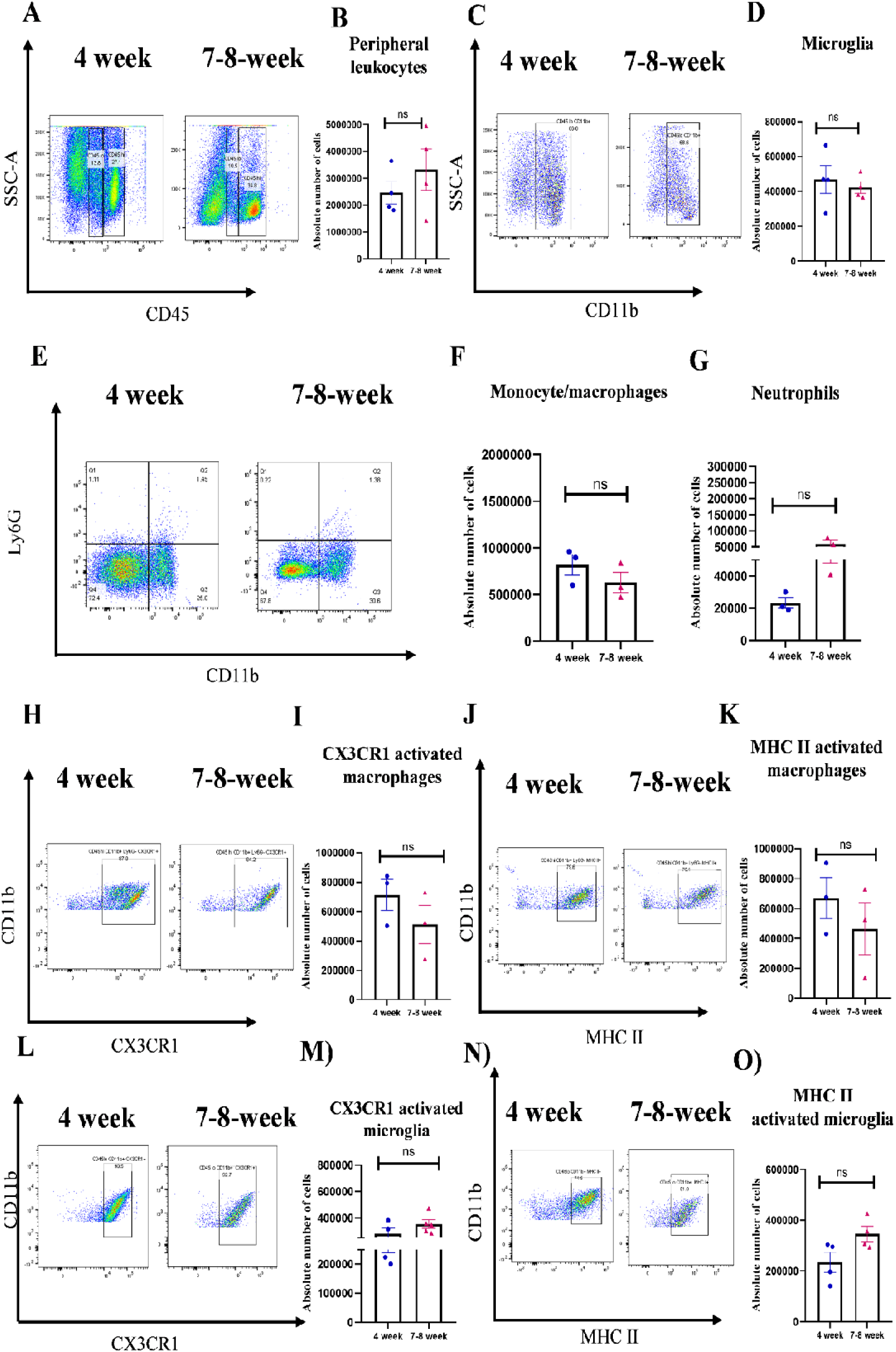
At the transition stage, the infiltration and activation of peripheral monocyte/macrophages, as well as the numbers and activation of brain-resident microglia, were comparable in mice from the two age groups. On day 7 p.i., brains from MHV-RSA59-infected 4-week and 7-8-week-old C57BL/6 mice were harvested for flow cytometry analysis and stained for CD45, CD11b, Ly6G, CX3CR1, and MHC II. Primary gating was performed on live cells, followed by singlets and CD45 hi. **A, C)** Representative flow cytometry dot plots showing percentages of overall CD45hi and CD45lo cell population and CD45lo CD11b+ (brain resident microglia). **B, D)** Graphical representation of the absolute cell numbers of peripheral leukocytes and brain resident microglia. **E)** Representative flow cytometry dot plots showing percentages of CD45hi CD11b+ Ly6G-(monocyte/macrophages) and CD45hi CD11b+ Ly6G+ (neutrophils). **F, G)** Graphical representation of the absolute cell numbers of monocyte/macrophages and neutrophils. Representative flow cytometry dot plots showing CD45hi CD11b+ Ly6G-cells expressing activation markers **H)** CX3CR1 **J)** MHC-II. Graphical representations of the absolute cell numbers have been shown in **I)** and **K),** respectively. Representative flow cytometry dot plots showing CD45lo CD11b+ cells expressing **L)** CX3CR1 **N)** MHC-II. Graphical representations of the absolute cell numbers have been shown in **M)** and **O),** respectively. Results were expressed as mean ± SEM. The shown flow cytometry study is representative of one independent experiment. (n=3-5 per age group) *Asterisk represents statistical significance calculated using an unpaired Student’s t-test; P<0.05 was considered significant.

